# Repeatome landscapes and cytogenetics of hortensias provide a framework to trace *Hydrangea* evolution and domestication

**DOI:** 10.1101/2024.06.05.597687

**Authors:** Shota Taniguchi, Sara Ishiguro, Nicola Schmidt, Matthias Jost, Stefan Wanke, Tony Heitkam, Nobuko Ohmido

## Abstract

**Background and Aims:** Ornamental hortensias are bred from a reservoir of over 200 species in the genus *Hydrangea* s.l. and are valued in gardens, households and landscapes across the globe. The phenotypic diversity of hortensia cultivars, hybrids and wild relatives is mirrored by their genomic variation, with differences in genome size, base chromosome numbers and ploidy level. We aim to understand the genomic and chromosomal basis of hortensia genome variation. Therefore, we analyze six hortensias with different origins and chromosomal setups for repeatome divergence, the genome fraction with the highest sequence turnover. This holds information from the hortensia’s evolutionary paths and can inform breeding initiatives.

**Methods:** We compiled a hortensia genotype panel representing members of the sections *Macrophyllae*, *Hydrangea, Asperae*, and *Heteromallae* and reconstructed a plastome-based phylogenetic hypothesis as evolutionary basis for all our analyses. We comprehensively characterized the repeatomes by whole genome sequencing and comparative repeat clustering. Major tandem repeats were localized by multi-color FISH.

**Key Results:** The *Hydrangea* species show differing repeat profiles reflecting their separation into the two major *Hydrangea* clades: Diploid *Hydrangea* species from Japan show a conserved repeat profile, distinguishing them from Japanese polyploids as well as Chinese and American hortensias. These results are in line with plastome-based phylogenies. The presence of specific repeats indicates that *H. paniculata* was not polyploidized directly from the common ancestor of Japanese *Hydrangea* species, but evolved from a distinct progenitor. Major satellite DNAs were detected over all *H. macrophylla* chromosomes.

**Conclusions:** Repeat composition among the *Hydrangea* species varies in congruence with their origins and phylogeny. Identified species-specific satDNAs may be used as cytogenetic markers to identify *Hydrangea* species and cultivars, and to infer parental species of old *Hydrangea* varieties. This repeatome and cytogenetics information helps to expand the genetic toolbox for tracing hortensia evolution and informing future hortensia breeding.

## Introduction

Hortensias are valued for their ornamental flowers (variable in color, size, and shape) as well as the growth form (from shrub to climber), but the underlying genomic and chromosomal variation is still not understood. Yet, this is needed to inform the many hortensia breeding initiatives around the globe (Wu and Alexander, 2020). The genus *Hydrangea* s.l. (Hydrangeceae), which contains more than 200 species and numerous subspecies (De Smet *et al*., 2015), is a group of woody flowering plants, likely originating from North America and East Asia (Raman *et al*., 2023; POWO, 2023). Molecular phylogenetic studies have shown the monophyly of *Hydrangea* s.l. with two main clades called *Hydrangea* clade I and II (Samain *et al*., 2010; Mendoza *et al*., 2013, 2014; De Smet *et al*., 2015; Raman *et al*., 2023). Several *Hydrangea* species native to Asia, including *H. macrophylla* and *H. serrata*, were brought to Europe about 200 years ago and since have been improved to serve as ornamental plants (Iwatsuki *et al*., 2001; Rinehart *et al*., 2006; Uemachi *et al*., 2014). During hortensia domestication, the genetic diversity of the cultivars decreased and beneficial traits have been lost (Uemachi *et al*., 2014). Therefore, it is important to assess the genome space and the relationships across hortensia wild relatives to inform local and global breeding programs.

The genetic landscape of *Hydrangea* is intricately shaped by variations in chromosome numbers and polyploidy levels, which exert profound effects on genome size, intraspecific diversity, and varietal identification. Notably, within the genus *Hydrangea*, polyploids can be present in one and the same species (e.g. *H. paniculata* individuals with chromosome numbers of 2n = 2x = 36, 3x = 54, 4x = 72, and 6x = 108; Funamoto and Tanaka, 1988). Furthermore, diploid *Hydrangea* species show variance in their chromosome number with 2n = 2x = 30-36 chromosomes (Table 1). These differences are due to chromosome rearrangements. Some *Hydrangea* species from Japan and China show aneuploidy, e.g. 2n = 2x = 30 in *H. involucrata*, and 2n = 2x = 34 in different *H. aspera* subspecies (Cerbah *et al*., 2001). Karyotype variations and genome size differences were not taken into account in the selection process during *Hydrangea* breeding, contributing to difficulties in the identification of the origin of certain *Hydrangea* varieties (e.g. *H. macrophylla* varieties).

**Table 1:**
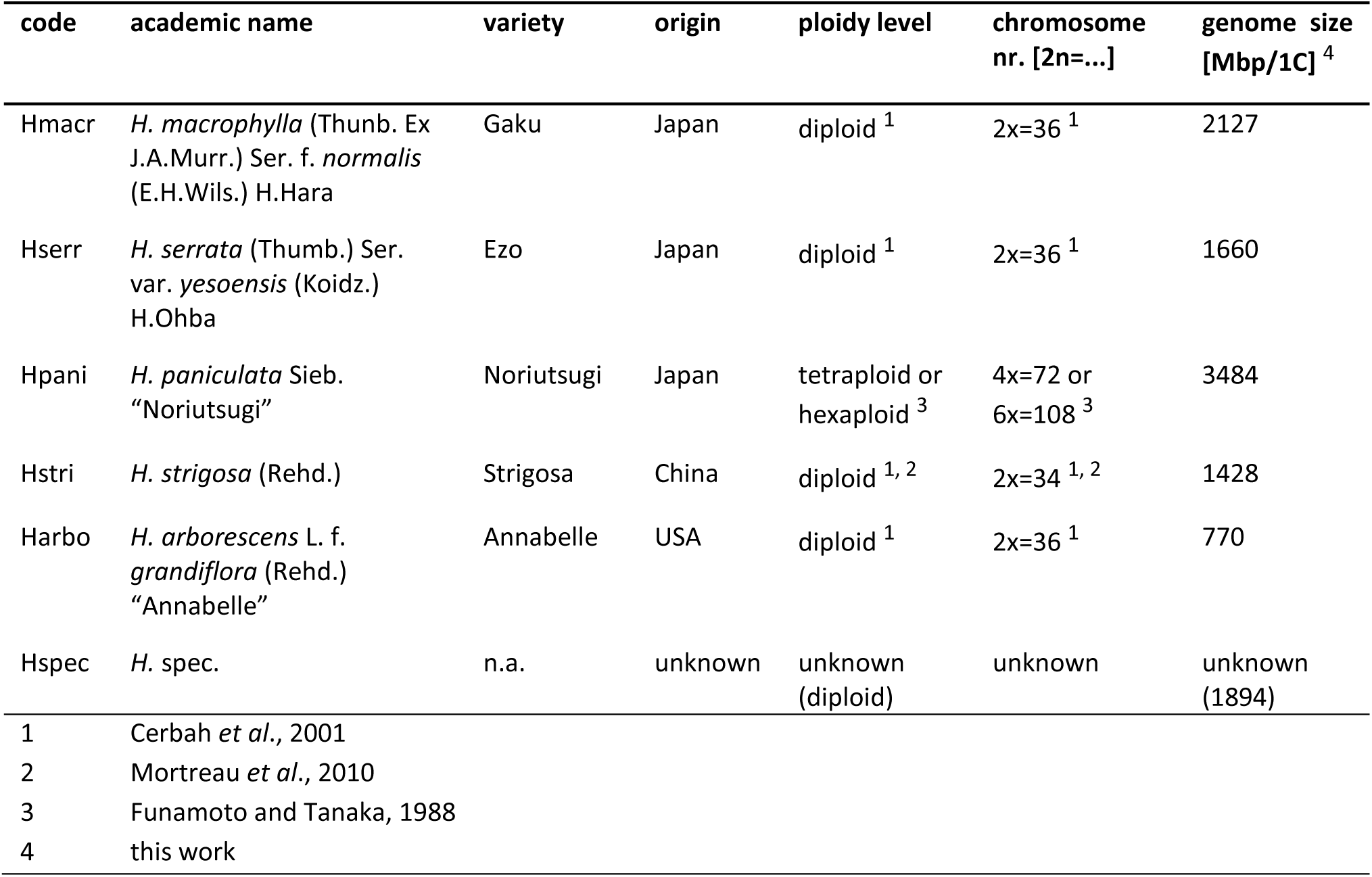
*Hydrangea* species sampled, including (suspected) ploidy level and genome size. In addition, a sixth undesignated genotype (*H*. spec.) was sampled, for which no further information (origin, chromosome number, genome size) is known. Since it is most closely related to *H. macrophylla* and *H. serrata* (see plastome-based phylogeny; Supplementary Data Figs. S1, S2), *H*. spec. is assumed to be diploid as well and its approximate genome size was calculated as mean value from the genome sizes of *H. macrophylla* and *H. serrata*.

Chromosomal mapping techniques, including fluorescence *in situ* hybridization (FISH), have the potential to offer this missing information (Heitkam and Garcia, 2023). By localizing individual repetitive sequences along the chromosomes, FISH can shed light onto the genomic dynamics underlying chromosome and karyotype evolution (Heslop-Harrison and Schmidt, 2012; Ohmido *et al*., 2007; Schmidt *et al*., 2019). For hortensias, cytogenetic methods using ribosomal DNA probes have already provided first insights into karyotype variation and the genomic intricacies among *Hydrangea* species (Van Laere *et al*., 2008; Mortreau *et al*., 2010). To understand the genetic basis of the chromosomal variation, and to develop probes for FISH-based cytogenetics, insights into the repetitive genome fraction of hortensias are needed.

Repetitive DNAs (repeats) build the structural backbone of the chromosomes and are the fastest-evolving sequences in a genome. Their fast sequence turnover and their genome-wide amplification and loss can drastically affect karyotypes and overall genome composition. The consequent genetic diversity potentially helps species adapting to environmental changes, thus leading to speciation (Paço *et al*., 2015). Thus, repeats often have species-specific genomic profiles. This helps to understand the relationships of species (Zagorski *et al*., 2020; Kuo *et al*., 2021) and to support taxonomic efforts (Reed and Rinehart, 2007, 2009; Chae *et al*., 2014; Dodsworth *et al*., 2015; Vitales *et al*., 2020). Depending on the repeat type, their analysis yields different information: Tandem repeats, such as satellite DNAs (satDNAs) and ribosomal DNAs (rDNAs), are often located at key chromosome positions. This includes the centromeres and telomeres, the ribosomal genes, and much of the constitutive heterochromatin (Biscotti *et al*., 2015). Hence, they are the prime choice to develop cytogenetic probes. In contrast, transposable elements (TEs) are dispersed in the genome and form different classes, lineages and families (Kubis *et al*., 1998; Sharma *et al*., 2005; Jurka *et al*., 2007). Their individual abundances and similarity profiles provide a fine-grained record of the evolutionary past of an organism (Heitkam *et al*., 2021; Schmidt *et al*., 2023).

To understand the hallmarks of *Hydrangea* genomes and to develop a framework for understanding the history of *Hydrangea* evolution and domestication, we here develop a 5-genotype-panel for *Hydrangea* (cyto-)genomics. This panel encompasses *Hydrangea* species from different geographic regions and reflects the genus-typical ploidy and base chromosome number variation. Here, we focus on *H. macrophylla* (Gaku), *H. serrata* (Ezo), *H. paniculata* (Noriutsugi), *H. strigosa* (Strigosa), and *H. arborescens* (Annabelle). To assess their suitability to serve as a genotype-panel for *Hydrangea* genomics, we determined their genome sizes in addition to previously published chromosome numbers and ploidies, and assigned an undesignated genotype (*H*. spec.) within this panel using plastome- and repeatome-based phylogenies. Repeatome analyses using low-coverage genome sequencing data was used to calculate the individual repeat fractions and to harness genomic differences between the genotypes. Finally, analyzing tandemly repeated sequences, we developed cytogenetic landmark probes with the aim to trace karyotype changes.

## Materials and Methods

### Plant material and genome size estimation

Genotypes of the genus *Hydrangea* were provided by the Kobe Municipal Arboretum (Kobe, Hyogo, Japan). The analyzed species encompass species belonging to both *Hydrangea* clades (I and II), and represent the following sections: *Macrophyllae* (*H. macrophylla* and *H. serrata*), *Hydrangea* (*H. arborescens*), *Asperae* (*H. strigosa*), and *Heteromallae* (*H. paniculata*) following the classification of De Smet *et al*. (2015). Additionally, an undesignated genotype (*H*. spec.) was included to determine its relationships to the above-mentioned species. Further accession details are provided in Table 1.

The six *Hydrangea* genotypes analyzed in this study represent small deciduous shrubs of two to five meters in diameter. *Hydrangea macrophylla* (“Gaku-ajisai” in Japanese; endemic to Japan) grows in Honshu and coastal forests (Iwatsuki *et al*., 2001). The inflorescence is composed of surrounding decorative petal-like-sepals and central non-decorative petals, corresponding to the lacecap inflorescence type (Fig. 1). Decorative sepals are blue to purple and white and non-decorative petals are blue to purple. *Hydrangea serrata* (“Ezo-ajisai” in Japanese; endemic to Japan) grows in northern Japan (Iwatsuki *et al*., 2001). The inflorescence resembles that of *H. macrophylla* (Fig. 1). *Hydrangea paniculata* (“Nori-utsugi” in Japanese), grows all over Japan (Iwatsuki *et al*., 2001). The inflorescence – a long hierarchical structure of white sepals and petals – is distinctly different from that of the two species mentioned above (Fig. 1). *Hydrangea strigosa* (“Strigosa” in Japanese) grows in China. The inflorescence resembles that of *H. macrophylla* (Fig. 1). However, decorative sepals are white. *Hydrangea arborescens* (“Annabel”) was propagated in the US. The inflorescence is composed of only decorative flowers corresponding to the mophead inflorescence type (Fig. 1).

**Fig. 1:**
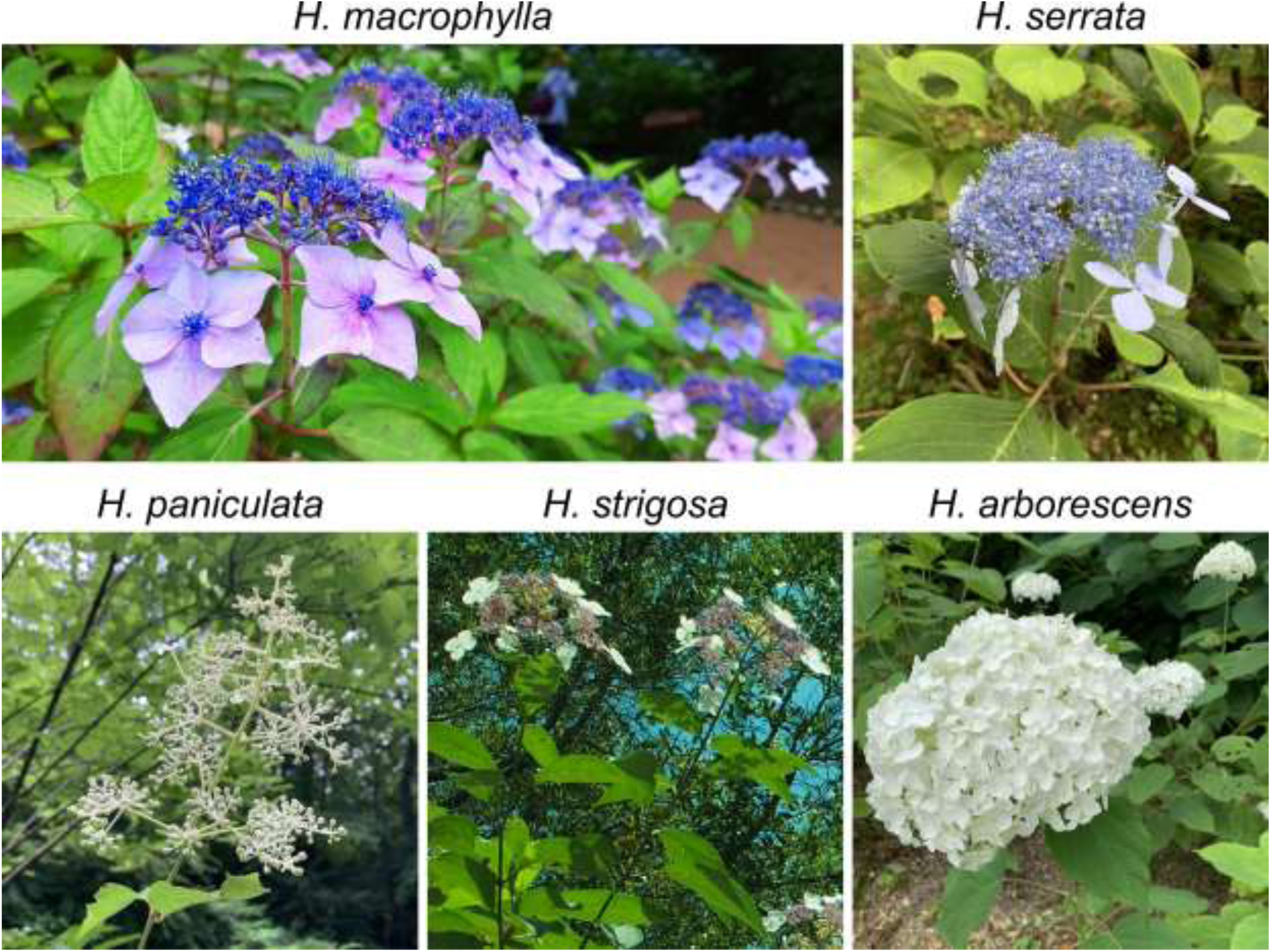
*Hydrangea* species used in this study. *Hydrangea macrophylla* grows in Japan mainland, Honshu (Pacific Ocean side of Kanto district), Izu Islands, Izu Peninsula, and Ogasawara Islands. Cutting cultivation is facilitated by its robust properties; leaves are large, thick and glossy. *Hydrangea serrata* grows in all the snowy northern areas of Hokkaido, Honshu (Japan, seaside from Aomori prefecture to Kyoto prefecture), and Kyushu (northern part and Osumi Peninsula). Leaves are rougher and thicker than those of *H. macrophylla*. *Hydrangea paniculata* grows in Hokkaido, Honshu, Shikoku, and Kyushu to Yakushima. The plant is the tallest of the Japanese hortensias. *Hydrangea strigosa* is native to central and southeast China. Its leaves are narrower and of a darker green compared to other *Hydrangea* species. *Hydrangea arborescens* is native to the Eastern United States. Its leaves are softer than those of the other *Hydrangea* species.

Flow cytometry was performed for the determination of genome size and ploidy level following Beckman Coulter’s application note with some modifications: The fresh leaf samples were immersed in 1.2 mL of chopping buffer (1.0 % Triton X-100, 140 mM 2-Mercaptoethanol, 50 mM NaHSO_3_, 50 mM Tris hydrochloric acid, 25 µg/mL propidium iodide). The leaves were then chopped with a new razor blade. The resulting nuclei suspension was passed through a 20 µm nylon mesh filter, placed on ice and another 0.6 mL of chopping buffer was added. The nuclei suspension was centrifuged at 1000 rpm for 4 min. The supernatant was then discarded and the pellet was resolved in another 0.5 mL of chopping buffer. The samples were analyzed using a flow cytometer (BD FACSAria TM III Cell Sorter) with measurements for each species taken on at least two different days. As an internal standard, *Oryza sativa* subsp. *japonica* “Nipponbare” (433 Mb/1C; Yu *et al*., 2005) was used.

### DNA isolation and sequencing

Genomic DNA was isolated from 10-100 mg lyophilized leaf material using the DNeasy Plant Mini Kit (QIAGEN, Hilden, Germany). DNA amount and quality were measured with the QubitTM 4 Fluorometer (Thermo Fisher Scientific, Inc., Waltham, MA, USA), and its integrity was checked via gel electrophoresis. Whole genome sequencing was performed to generate 2×150 bp paired-end (PE) reads with the Novaseq 6000 system (Illumina, San Diego, CA, USA) by AZENTA, formerly Genewiz.

### Plastome reconstruction for phylogenetic tree estimation

By sequencing a series of chloroplast regions and internal transcribed spacer (ITS) regions for an extensive sampling of *Hydrangea* specimen, phylogenetic constructions of the tribe Hydrangeae have been performed (Samain *et al*., 2010; De Smet *et al*., 2015; Raman *et al*., 2023). In this study, we reconstructed the plastid genomes of the newly sequenced genotypes, and used them for phylogenetic tree reconstruction as an independent evidence for comparison with ploidy levels and nuclear repeat content as well as positioning the undesignated *Hydrangea* species in the taxonomic framework.

The raw data of the whole sequences were trimmed for adapters and quality with Trimmomatic (v0.39; Bolger *et al*., 2014) using standard settings. The read quality was visualized with FastQC (v0.11.9; Andrews, 2010) before and after trimming. A gold standard for assembly and annotation of high-quality plastomes based on *de novo* assembly methods and appropriate references for gene annotation was used (Jost and Wanke 2024). *De novo* plastome assemblies were generated using GetOrganelle (v1.7.7.0; Jin *et al*., 2020), setting the number of rounds to 50. The resulting scaffolds were imported into Geneious (v11.1.5) and finalized to create circular plastomes, where necessary. Correctness of the assemblies was verified via red mappings using bwa (Li & Durbin, 2009) and by alignments to published *Hydrangea* data. Genes were annotated using the plastome of *Jamesia americana* (NC_044836; Fu *et al*., 2019) as reference. The 5’-end of the *atp*B reference gene was moved to the next canonical in-frame start codon, resulting in a 3 bp shortening of the sequence. The reference for the plastid *ycf*15 gene was taken from the plastome of *Cornus bretschneideri* (MN651479; Li *et al*., 2020), as this gene was not annotated on the *Jamesia* (NC_044836) plastome. TransferRNA boundaries were additionally verified using tRNAscan-SE (Lowe and Chan, 2016). Two different data sets were analyzed for their phylogenetic signal, namely the complete plastome (excluding one inverted repeat copy) and the protein-coding and rRNA gene information only. The outgroup sampling is comprised of representatives from the sister tribe Philadelpheae (*Kirengeshoma palmata* NC_044808; Fu *et al*., 2019) and the sister subfamily Jamesioideae (*Jamesia americana* NC_044836; Fu *et al*., 2019). *Eucnide grandiflora* (NC_044767; Fu *et al*., 2019), a member of sister family Loasaceae, was used for rooting. The alignments were generated using MAFFT (v7.450; Katoh *et al*., 2002; Katoh and Standley, 2013) and manually adjusted in AliView (v1.28; Larsson, 2014). The best-fit nucleotide substitution models for multiple sets of partitions and data were estimated using ModelFinder (Kalyaanamoorthy *et al*., 2017) and used as input for tree reconstruction using both RAxML (allowing only for the GTR model; Stamatakis, 2014) and IQ-TREE (Nguyen *et al*., 2015). Support values are based on 1,000 bootstrap replicates. Inferences were then visualized using TreeGraph2 (Stöver and Müller, 2010).

### Read pre-processing and RepeatExplorer2 clustering

After the quality assessment (see section above), raw Illumina reads were trimmed to a final length of 100 nt (20 nt were trimmed from the 5’ ends and 30 nt from the 3’ ends). Trimmed reads were randomly sampled to obtain 0.1× and 0.05× genome coverage and converted to fasta format using Seqtk (https://github.com/lh3/seqtk). Since no genome size was known for the undesignated genotype *H*. spec., an approximate genome size was assumed based on the plastome-derived phylogeny: *H*. spec. proved to be the closest relative to *H. macrocarpa* and *H. serrata* (Supplementary Data Figs. S1, S2). Thus, the genome size of *H*. spec. was calculated as the mean value of the genome sizes from *H. macrocarpa* and *H. serrata* (1894 Mbp/1C). A graph-based clustering method was performed to identify, characterize and quantify repetitive sequences in each *Hydrangea* genome using the RepeatExplorer2 pipeline on the Galaxy server (https://repeatexplorer-elixir.cerit-sc.cz) with default settings (Novák *et al*., 2010, 2013). The automatic repeat annotation was manually revised. Repeat abundances were individually assessed based on the read amount within the respective cluster for each species (0.1× genome coverage). Additionally, a comparative read clustering was performed with sampled PE reads equivalent to 0.1× or 0.05× genome coverage based on the respective ploidy level (Table 1; *H*. spec. was assumed to be diploid). Since the analyzed *H. paniculata* individual is assumed to be tetraploid (see Discussion), a 0.05× genome coverage was used to match the genome size of the remaining diploid individuals (0.1× genome coverage). Each PE read dataset was labeled with a unique five letter code (Table 1). The comparative clustering was performed with default settings.

### Bioinformatic repeat characterization

Candidate species-specific and conserved satDNAs were identified based on the comparative RepeatExplorer2 analysis. A BLAST search of the consensus monomer sequences against *H. macrophylla* long reads was performed to confirm their arrangement in long tandem arrays. Although genomic resources for *Hydrangea* are not abundant, long assembled genome sequences in *H. macrophylla* ‘Aogashima-1’ are available (Nashima *et al*., 2021). Fourteen long read datasets (DRX222164) were downloaded from the European Nucleotide Archive. Monomeric sequences were aligned to the long reads using blastn (Camacho *et al*., 2009), and the long reads containing BLAST hits were assessed with self-dotplots that were constructed using FlexiDot (Seibt *et al*., 2018).

### Probe generation

Amplification of *Hydrangea* satDNA fragments was performed with specific primer pairs designed based on the RepeatExplorer2-derived satDNA monomer consensus sequences (Supplementary Data S1). Detailed information on the primers can be found in the Supplementary Data Table S1. The target sequences were amplified using standard PCR reactions with genomic DNA of *H. macrophylla*. An initial denaturation at 94 °C for 3 min was followed by 35 cycles of denaturation at 94 °C for 45 s, primer-specific annealing temperature for 30 s and extension at 72 °C for fragment-specific extension time. A final extension at 72 °C for 5 min was performed. The PCR fragments were purified, cloned and commercially sequenced. Sequenced inserts with the highest identity to the consensus monomer sequences, comprising two up to three complete repetitions of the respective monomer, were used as probes for the following hybridization experiments (Supplementary Data S2).

### Chromosome preparation and FISH

Mitotic chromosomes were prepared from actively growing meristems using young buds (bud diameter: 0.7 up to 1.7 mm) from *H. macrophylla*. The buds were fixed in methanol:glacial acetic acid (3:1) and stored at −20 °C until required. For sample preparation, a bud was washed in water thoroughly, digested in enzymatic maceration containing 2.5% Pectolyase Y-23 (Seishin Pharmaceutical Co., Ltd, Chiba, Japan) and 1% Cellulase Onozuka RS (Yakult Honsha Co., Ltd, Tokyo, Japan), and vacuumed for 5 min in a desiccator under −0.1 mPA and incubated at 37 ℃ for 60 min. The bud was then macerated in a few drops of ethanol:acetic acid (3:1) using fine forceps following the air-drying method (Ohmido *et al*., 1998).

The FISH experiments were performed using the protocol of Schmidt *et al*. (1994) and Liu *et al*. (2021). The probe “18SrRNAgene_Bv_probe1” (Sielemann *et al*., 2023) was used for the detection of the rDNA (directly labeled with DY415). Specific *Hydrangea* satDNA probes were labeled indirectly by PCR with biotin-16-dUTP (Roche Diagnostics) detected by streptavidin-Cy3 (Sigma–Aldrich). Chromosome slides were counterstained with 2 µg/mL 4’,6-Diamidino-2-phenylindole (DAPI). FISH images were photographed directly using a ASI BV300-20A camera coupled to a Zeiss Axioplan 2 imaging microscope with appropriate filters. Image analyses were performed using the Applied Spectral Imaging v. 3.3 software (ASI).

### Access to data

Illumina whole genome sequence data are available at EBI under the accession numbers ERR12526782–ERR12526787 (Study ID: ERP151402; Supplementary Data Table S2). The complete plastome sequences of the six *Hydrangea* individuals analyzed in this study have been deposited in GenBank under the accession numbers OR701877-OR701882 (Supplementary Data Table S2). Satellite DNA consensus sequences and the sequences used as FISH probes are available in the Supplementary Data S1 and Data S2. Furthermore, cloned sequences of the satDNAs *Hydrangea*TR01-05a, 07, and 13-15 were submitted to ENA (accession numbers OY742023-OY742031).

## Results

### A genotype-panel to characterize the genus *Hydrangea*

The *Hydrangea* species investigated in this study were selected to span remarkable differences in hortensia chromosome numbers, both in terms of base chromosome number and ploidy level (Table 1). Whereas *H. macrophylla*, *H. serrata*, and *H. arborescens* represent diploids with 2n = 2x = 36 chromosomes, the diploid chromosome set of *H. strigosa* comprises only 2n = 2x = 34 chromosomes. On the other hand, *H. paniculata* includes diploid, triploid, tetraploid as well as hexaploid plants with chromosome numbers of 2n = 36, 54, 72, and 108, respectively (Funamoto and Tanaka, 1988).

Although more genomic data of hortensias became available in the last decade, most of the available phylogenies include only a fraction of the large *Hydrangea* diversity. As a result, the *Hydrangea* systematics is still in flux. Exploring the possibility of using the repeatome to classify a species without genetic information, one undesignated species from the Kobe Municipal Arboretum (*H*. spec.) was included in our genotype-panel. We derived a phylogeny of all genotypes used in this study based on their plastomes (protein-coding and rRNA regions as well as whole plastid genomes; Supplementary Data Figs. S1, S2). Plastome reconstruction resulted in complete, circular genomes for all accessions. The plastid chromosomes possess the typical quadripartite structure and range from 157,617 bp to 158,055 bp in size (Supplementary Data Fig. S1). The reconstructions share identical gene and intron content. The tree reconstructions (Supplementary Data Fig. S2), depict extremely short branch lengths for *H. macrophylla*, *H. serrata*, and *H*. spec. being the likely reason for varying topologies, depending on the data set used (bootstrap support values between 52-81; Supplementary Data Fig. S2) and pointing to a very close relationship of *H*. spec. to the two members of *Hydrangea* clade II, thus, also being a member of the section *Macrophyllae*.

Similar to ongoing phylogenetic investigations, information on *Hydrangea* genome sizes are patchy and sometimes vary considerably ([Plant DNA C-values Database](https://cvalues.science.kew.org/)). Therefore, we estimated the genome sizes for the *Hydrangea* genotypes designated to a certain species (Table 1). The genome sizes among these five *Hydrangea* species range from 770±83 Mb/1C in *H. arborescens* to 3484±52 Mb/1C in *H. paniculata*. Among the Japanese diploid *Hydrangea* species, the genome size of *H. macrophylla* (2127±273 Mb/1C) is larger than that of *H. serrata* (1660±108 Mb/1C) although their chromosome numbers are the same. In contrast, *H. strigosa* shows both a reduced chromosome number (34 chromosomes) and a lower genome size (1428±94 Mb/1C). The observed variation in origin, chromosome number, and genome size indicates that the *Hydrangea* species of this study represent a powerful genotype panel for the analysis of the *Hydrangea* diversity.

### *Hydrangea* genomes are mainly composed of Ty3/Gypsy retrotransposons

To estimate and compare the repeat compositions among the six *Hydrangea* genotypes, clustering analyses were individually performed with the RepeatExplorer2 pipeline. The number of reads per sample was between 0.77-3.48 million, corresponding to a 0.1× genome coverage of the respective species (Table 1). Based on the threshold of 0.01% (minimum number of reads within a cluster; at least 71-348 reads), 338 up to 403 clusters were annotated as repetitive DNA, accumulating in 204 up to 260 superclusters. This equals an overall repeat proportion of 42.15 up to 61.08%, highly correlating with the genome size (r^2^ = 0.99; Fig. 2; Table 2).

**Fig. 2:**
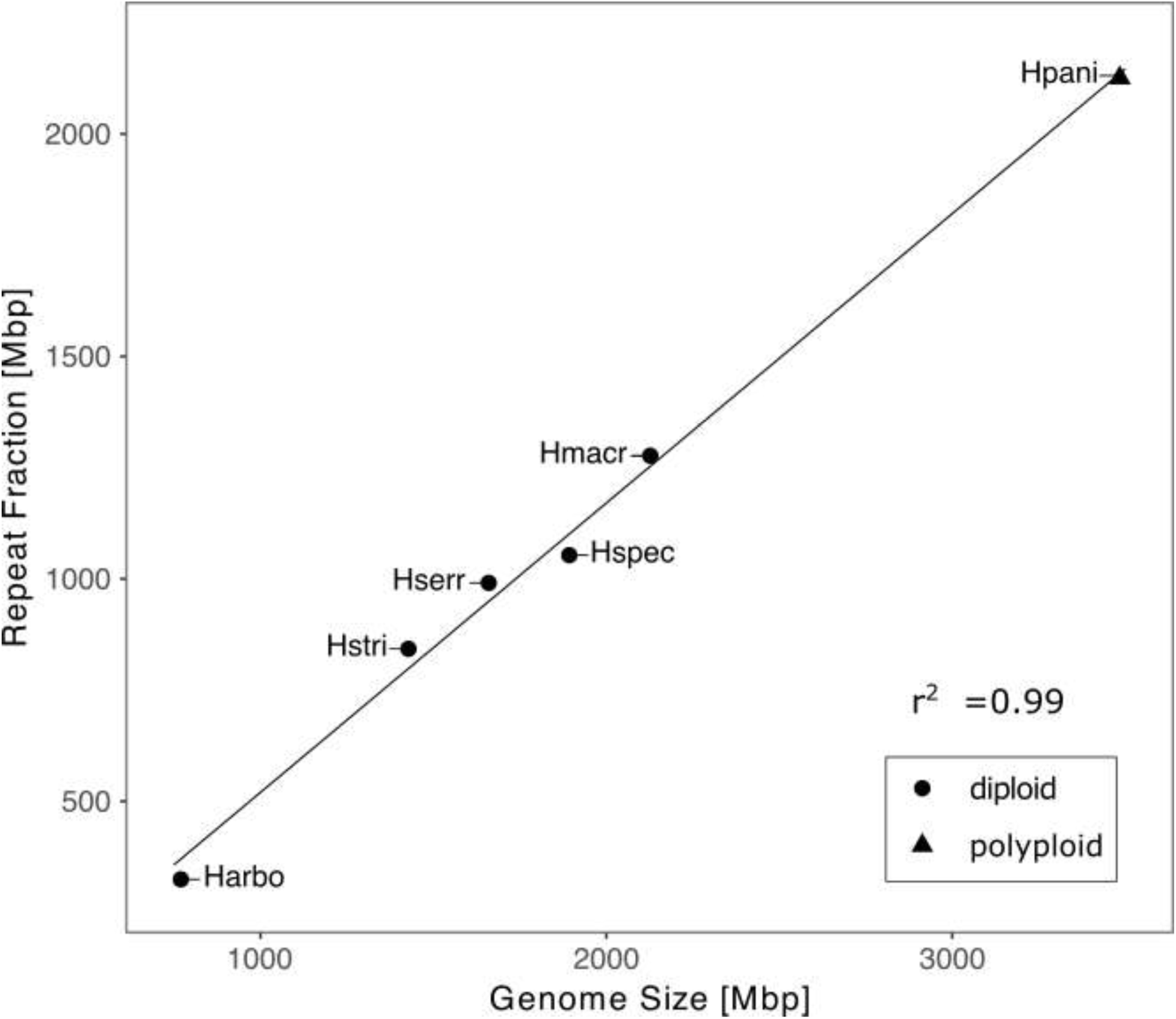
Correlation between genome size and repeat fraction among six different *Hydrangea* genotypes. Shapes indicate different ploidy levels. The sample abbreviations are according to Table 1.

**Table 2:**
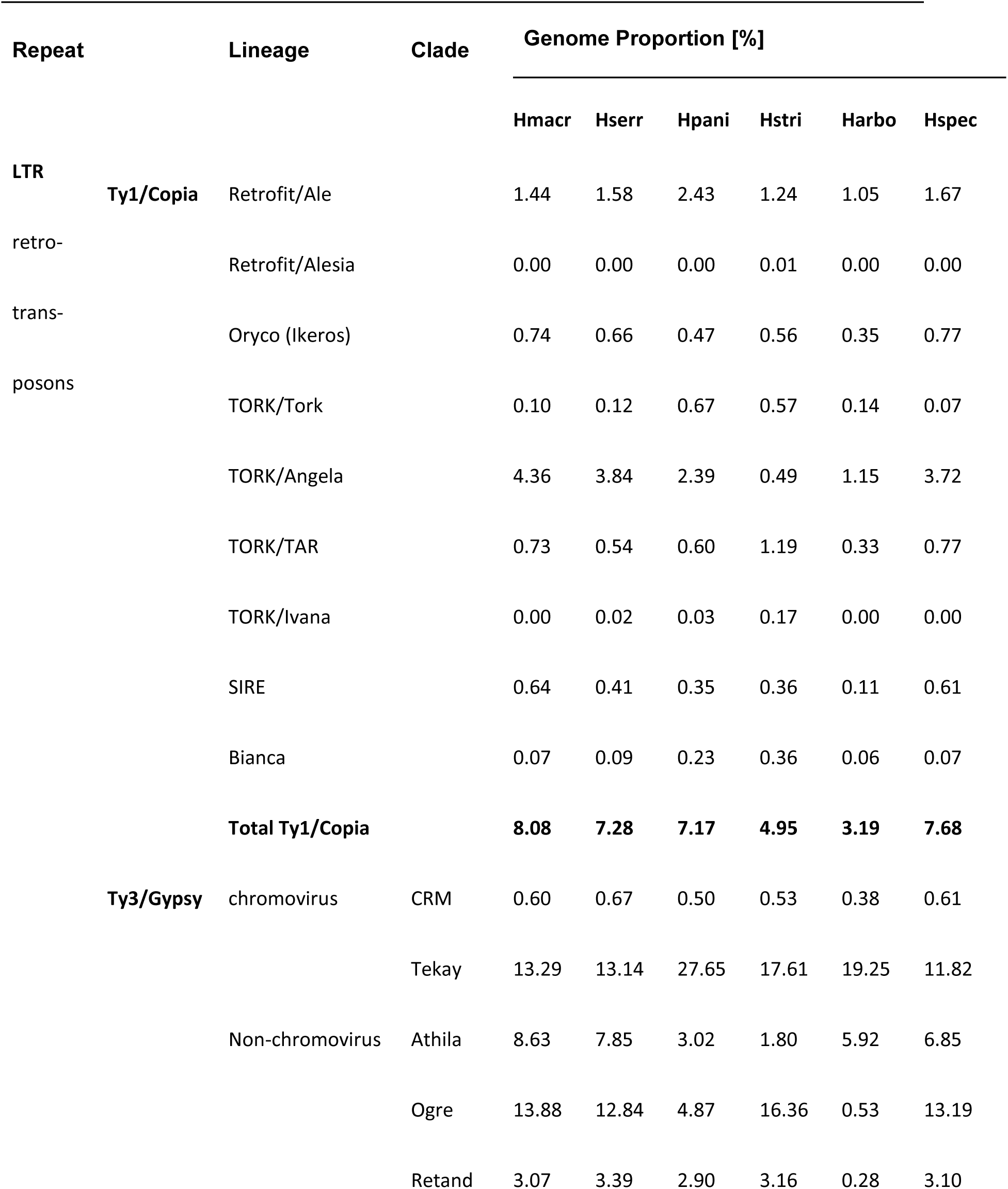
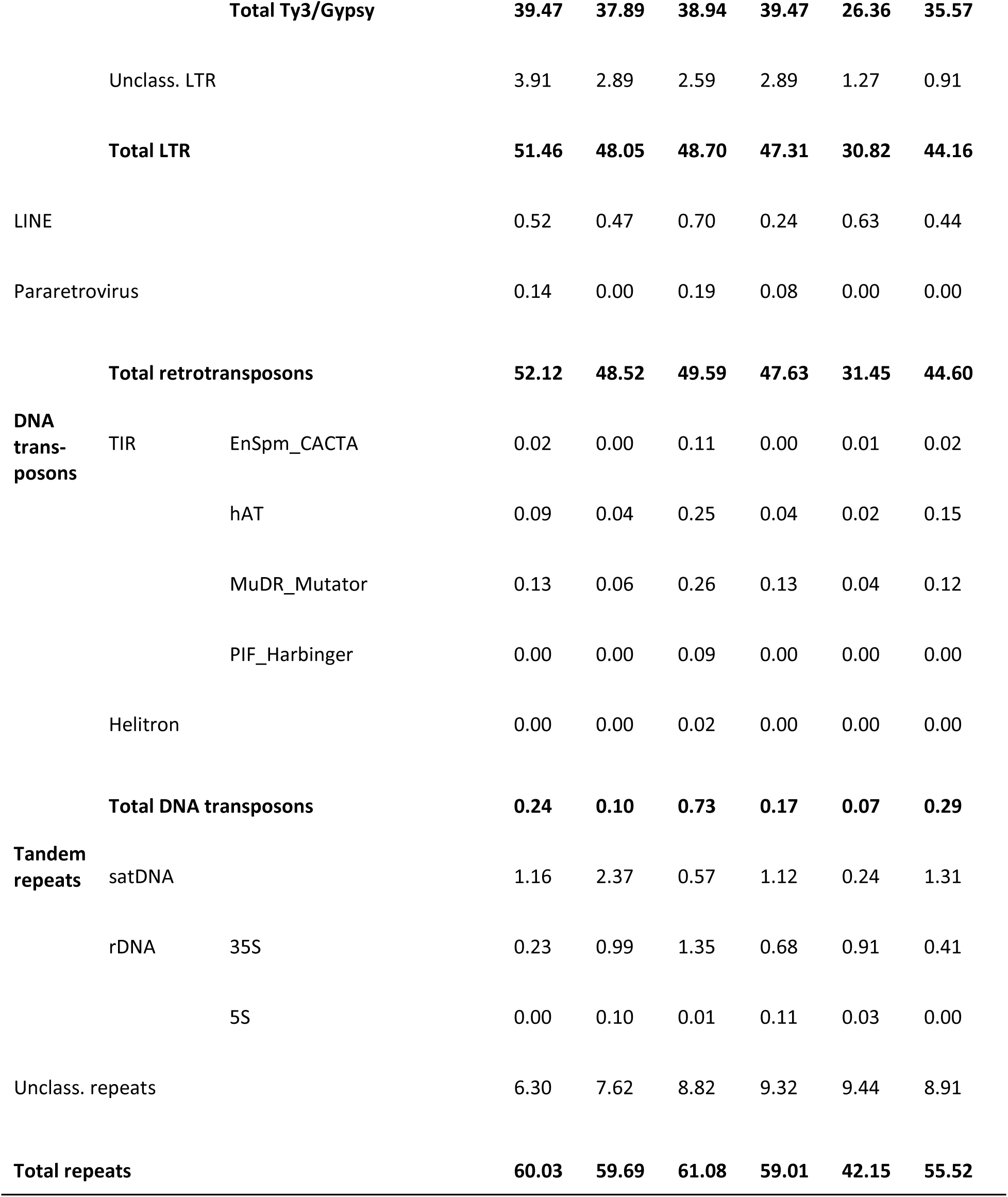
Genome proportions [%] of different repeat classes among six *Hydrangea* genotypes (abbreviations as in Table 1). The corresponding absolute repeat amounts ([Mbp] based on the genome sizes listed in Table 1) are shown in the Supplementary Data Table S3.

The LTR retrotransposons are the most abundant repetitive elements within the *Hydrangea* genomes accounting for 30.82 up to 51.46% (Fig. 3; Table 2). Retrotransposons from the Ty3/Gypsy superfamily are the predominant elements within this fraction (26.36-39.47%), represented by four clades; Tekay and CRM from the chromovirus lineage, as well as Athila and Tat (subclades Ogre and Retand) from the non-chromovirus lineage (Fig. 4). Tekay elements stand out showing genome proportions of 11.82% up to 27.65%. Ogre elements show the highest variety in abundance with only 0.53% in *H. arborescens* compared to 16.36% in *H. strigosa* (Table 2). However, Athila elements also show variance in abundance as they occupy 5.92-8.63% of the genomes from *H. macrophylla*, *H. serrata*, *H. arborescens*, and *H*. spec., whereas they only account for 1.80% and 3.02% in *H. strigosa* and *H. paniculata*, respectively. Retand elements show a species-specific low abundance in *H. arborescens* with a genome proportion that is less than one-tenths compared to that of the other *Hydrangea* species (Fig. 4).

**Fig. 3:**
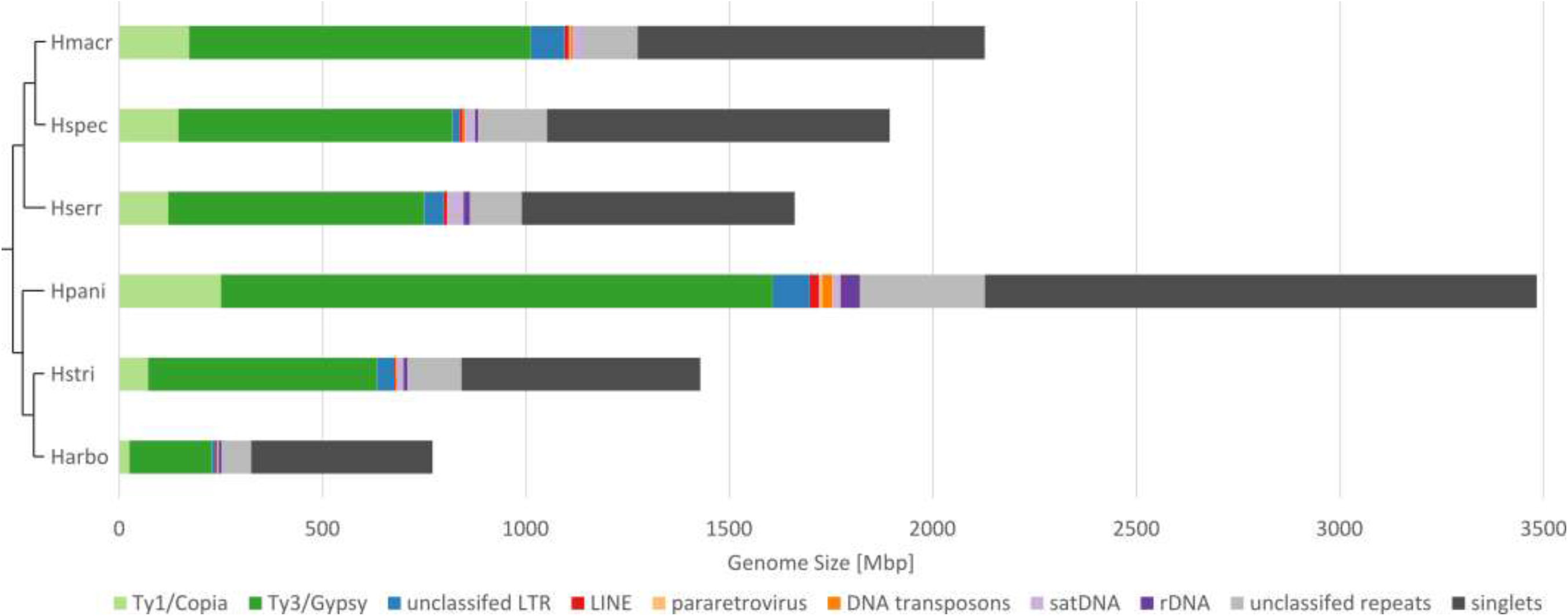
Repeat composition of the six *Hydrangea* genotypes. The sample abbreviations are according to Table 1. The displayed cladogram is based on the plastome-derived phylogeny (protein-coding and rRNA regions of the plastid genome; see Supplemental Data Fig. S2). The proportion of major repeat classes is shown in relation to the total genome size.

**Fig. 4:**
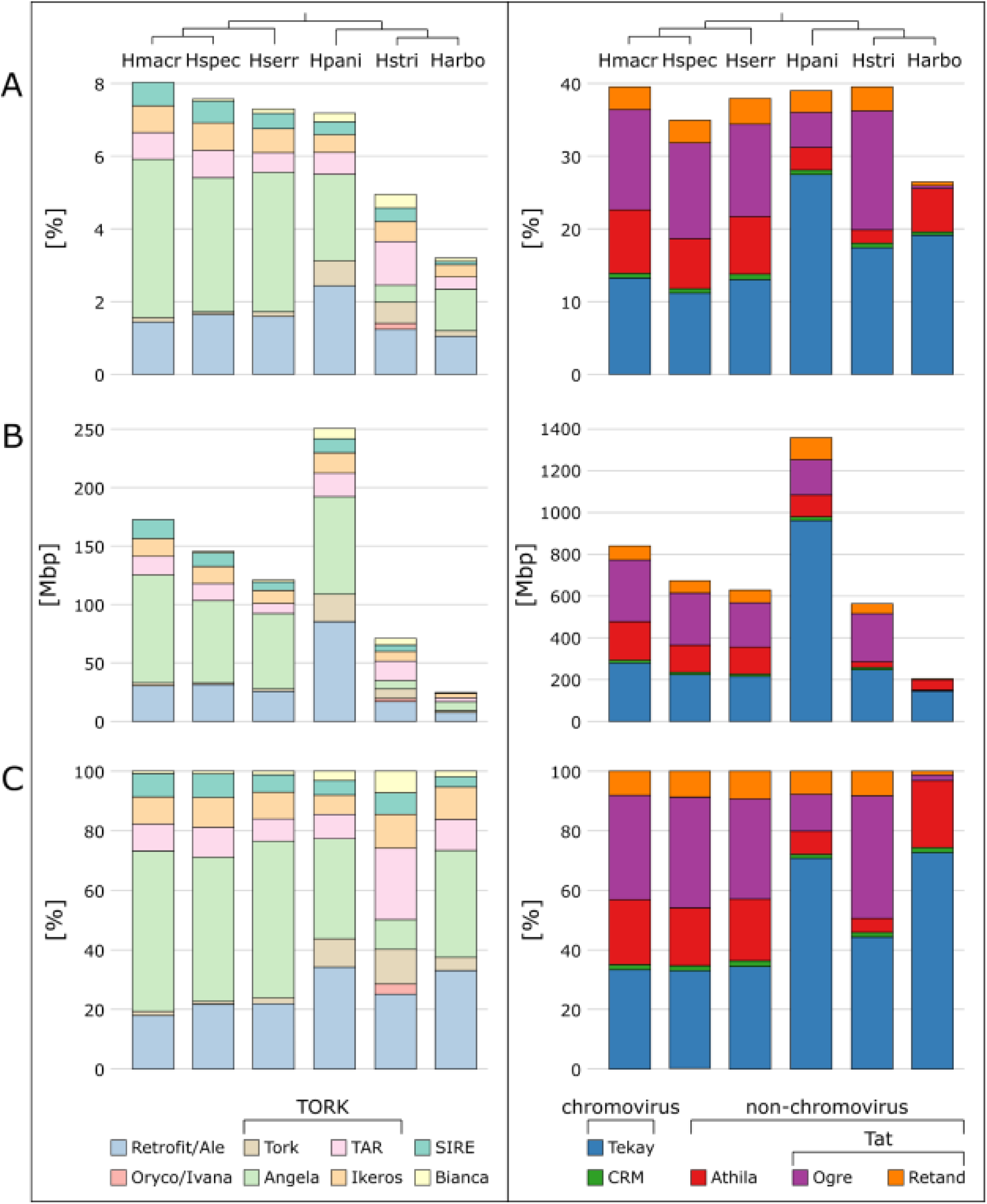
Repeat composition of LTR retrotransposons (left: Ty1/Copia; right: Ty3/Gypsy) among the six *Hydrangea* genotypes. The sample abbreviations are according to Table 1. The displayed cladogram is based on the plastome-derived phylogeny (protein-coding and rRNA regions of the plastid genome; see Supplemental Data Fig. S2). (A) Percentage contribution of the respective Ty1/Copia (sub-)lineage or Ty3/Gypsy clade to the *Hydrangea* genomes. Please note the different scales. (B) Contribution of the respective Ty1/Copia (sub-)lineage or Ty3/Gypsy clade to the *Hydrangea* genomes as absolute values [Mbp]. Please note the different scales. (C) Relative composition of the LTR retrotransposon fractions.

Ty1/Copia retrotransposons account for 3.19% up to 8.08% of the *Hydrangea* genomes (Fig. 3; Table 2), being composed of five lineages; Retrofit (clades Ale, Alesia), Oryco/Ivana, TORK (clades Tork, Angela, TAR, and Ikeros), Bianca, and SIRE (Fig. 4). Among diploid *Hydrangea* species with Japanese origin (*H. macrophylla* and *H. serrata*), Angela elements are predominant, occupying 3.84-4.36% of the respective genome. On the other hand, Angela and Ale elements are similarly abundant in the genomes of *H. paniculata* (approx. 2.5%, each) and *H. arborescens* (approx. 1%, each), whereas Ale elements are the most abundant Ty1/Copia retrotransposons in *H. strigosa* (1.24% of the genome; Table 2). DNA transposons contribute 0.07-0.73% to the *Hydrangea* genomes (Table 2). In regards to the tandem repeats, satDNAs occupy 0.24 up to 2.37% of the respective genome.

### Repeat composition patterns distinguish Chinese and American from Japanese *Hydrangea* species

To compare the repeat composition among the six *Hydrangea* genotypes and to identify species-specific repetitive sequences, comparative read clustering was performed. The number of reads per sample ranged from 0.77 to 2.14 million (a total of 9.869.000 reads), corresponding to a genome coverage of 0.1× for the diploid species and 0.05× for the assumed tetraploid *H. paniculata*. The RepeatExplorer2 pipeline reached an analysis limit by retrieving 8.492.761 reads. The total repeat proportion was calculated as 62.82% (represented by 447 clusters within 302 superclusters; based on the minimum read number threshold of 0.01% per cluster). Consequently, 37.18% of the reads were considered non-repetitive.

The six *Hydrangea* genotypes analyzed were divided into four groups based on the composition and abundance of identified repeats in their genomes; Japanese diploid *Hydrangea* (*H. macrophylla*, *H. serrata*, and *H*. spec.), Japanese polyploid *Hydrangea* (*H. paniculata*), Chinese *Hydrangea* (*H. strigosa*), and American *Hydrangea* (*H. arborescens*; Fig. 5). Whereas *H. macrophylla*, *H. serrata*, and *H*. spec. share nearly all clusters, indicating a highly similar repeat composition among these three genotypes, the genomes of *H. paniculata*, *H. strigosa*, and *H. arborescens* each show unique repeat patterns.

**Fig. 5:**
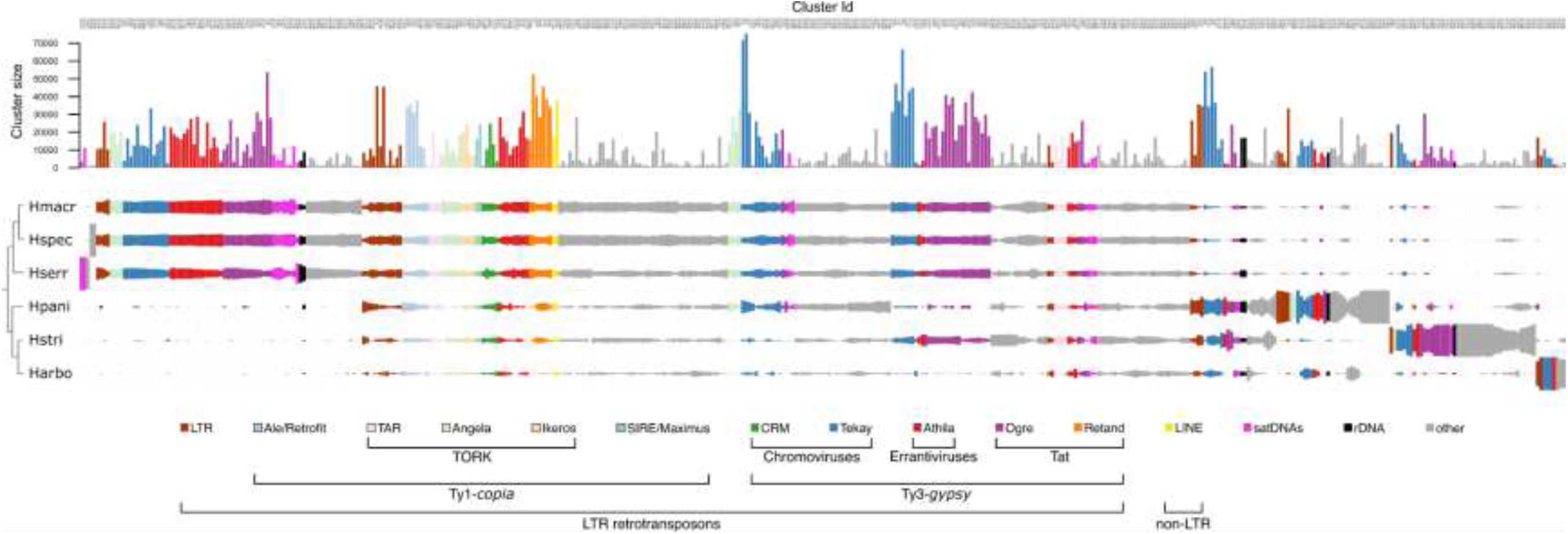
Comparative repeat composition among *Hydrangea* species. The bars represent the distribution of clusters comprising at least 850 reads ( ≥ 0.01% of the analyzed reads) among the analyzed species. Rectangles are colored according to the type of repetitive element and their size is proportional to the genomic abundance in the respective species (abbreviations as in Table 1). The displayed cladogram is based on the plastome-derived phylogeny (protein-coding and rRNA regions of the plastid genome; see Supplemental Data Fig. S2).

The genome of the polyploid *H. paniculat*a, which is endemic to Japan, has the largest overall repeat fraction among the analyzed species (more than 61%; see Table 2). Furthermore, it harbors several specific LTR retrotransposons, including Angela, Athila, and Ogre members (one each) and three different Tekay elements. The similarly sized repeat fraction of the *H. strigosa* genome (more than 59%; see Table 2) includes two specific Ogre elements. The *H. arborescens* genome has the smallest total repeat fraction among the species analyzed (less than 43%; see Table 2), but also contains specific LTR retrotransposons, one of which is a Tekay element. Although the overall repeat composition of the *H. serrata* genome resembles those of *H. macrophylla* and *H*. spec., two *H. serrata*-specific satellite DNAs were identified within its genome.

### Hallmarks of *Hydrangea* tandem repeats

As tandem repeats often span large genomic regions, they are prime targets for the development of cytogenetic landmark probes. In addition, as they are among the most rapidly evolving genome components, they have potential as genome-specific probes. Hence, we focused on the most abundant *Hydrangea*-specific tandem repeats, delimiting their nucleotide sequence, abundance, degree of repetitiveness and their potential to serve as cytogenetic probes.

The comparative clustering analysis revealed 16 different satDNAs with species-specific abundances among the six *Hydrangea* genotypes (Fig. 6). These satDNAs were named “*Hydrangea* tandem repeat” (*Hydrangea*TR), and numbered based on their overall abundance throughout the six *Hydrangea* genomes. As two monomer variants were identified for *Hydrangea*TR05, the variants with the overall higher and lower abundances were called *Hydrangea*TR05a and *Hydrangea*TR05b, respectively. *Hydrangea*TR05a is abundant in the Japanese *Hydrangea* species including the polyploid *H. paniculata*. However, its variant *Hydrangea*TR05b is enriched in the *H. paniculata* as well as in the American *H. arborescens* genome (Fig. 6). Some HydrangeaTRs are enriched in certain *Hydrangea* species, such as *Hydrangea*TR04 that is comparatively abundant in the genomes of *H. paniculata* and *H. strigosa*. Species-specific satDNAs were identified for *H. serrata* (*Hydrangea*TR02 and 10) as well as for *H. paniculata* (*Hydrangea*TR13).

**Fig. 6:**
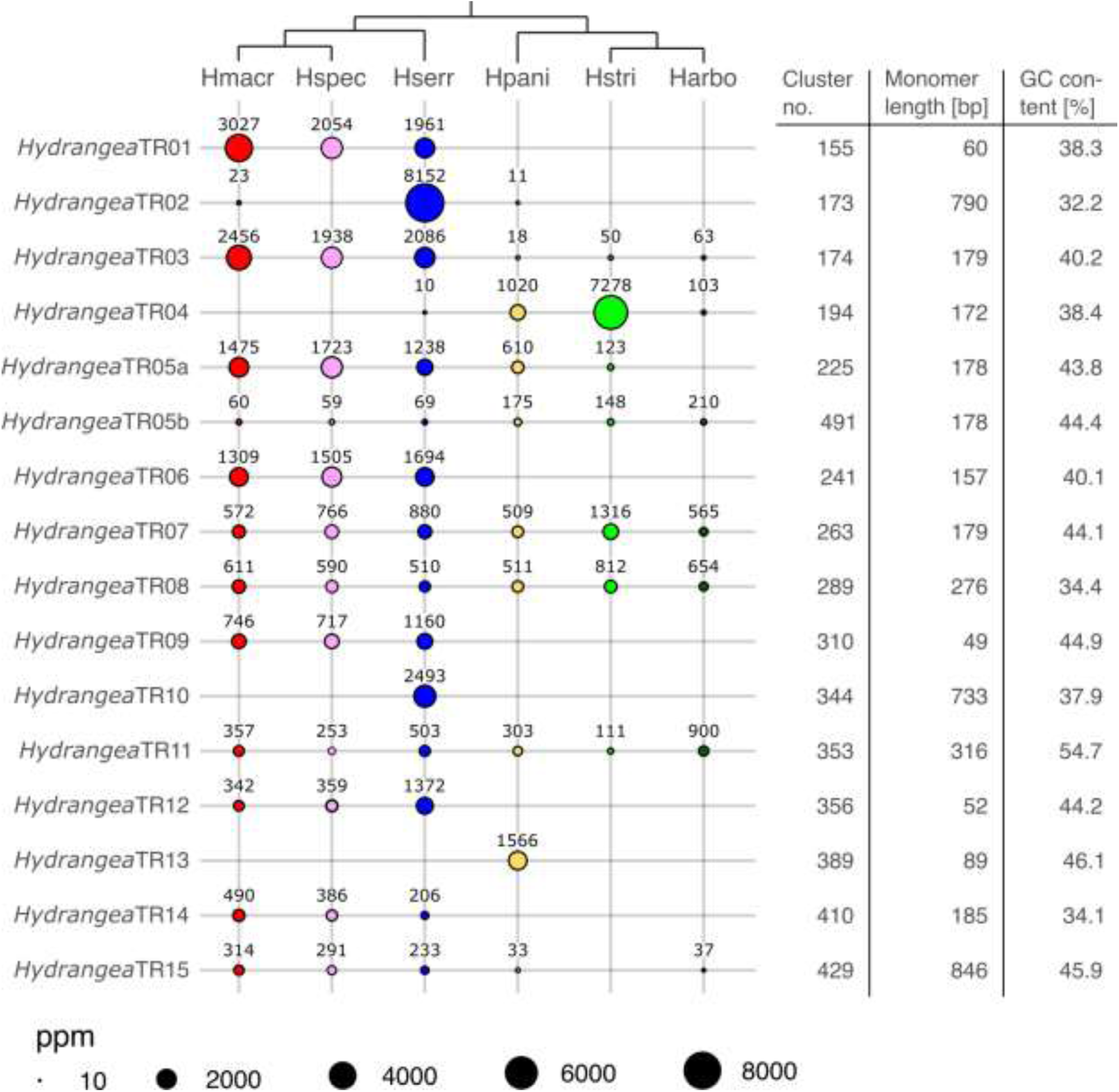
Quantification of satDNAs within different *Hydrangea* genomes and further *Hydrangea*TR characteristics. The relative satDNA abundance is indicated by the size of the circles, which corresponds to the genome proportion as parts per million [ppm]. The sample abbreviations are according to Table 1. The displayed cladogram is based on the plastome-derived phylogeny (protein-coding and rRNA regions of the plastid genome; see Supplemental Data Fig. S2).

To determine whether the HydrangeaTRs have the capacity to form long tandem arrays and thus represent canonical satDNAs, we investigated their genomic environment on long reads: The monomer sequences of the 16 identified *Hydrangea*TRs were mapped to the long read dataset of *H. macrophylla* (GenBank accession number DRR231909; Nashima *et al*., 2021) and the continuity of the arrays was estimated visually using self dotplots (Supplementary Data Fig. S3). Arrangement into continuous arrays could be verified for the twelve *Hydrangea*TRs that occur in the *H. macrophylla* genome, with the remaining *Hydrangea*TRs being restricted to other *Hydrangea* species. Therefore, the detection of long tandem arrangements of *Hydrangea*TR02, 04, 10, and 13 was not possible in *H. macrophylla* (Supplementary Data Fig. S3).

### Chromosomal localization of tandem repeats along *Hortensia* chromosomes

Based on the initial characterization of *Hydrangea* satDNAs, we selected the three most abundant tandem repeats of the *H. macrophylla* genome for fluorescent *in situ* hybridization; *Hydrangea*TR01, 03, and 05a. To determine the chromosomal localization of these satDNAs, mitotic chromosomes of *H. macrophylla* (2n=2x=36) were hybridized with biotin-labeled probes marking *Hydrangea*TR01, 03, and 05a, respectively (Fig. 7, red signals), together with a probe marking the 18S rDNA as part of the 35S rDNA (Fig. 7, turquoise signals).

**Fig. 7:**
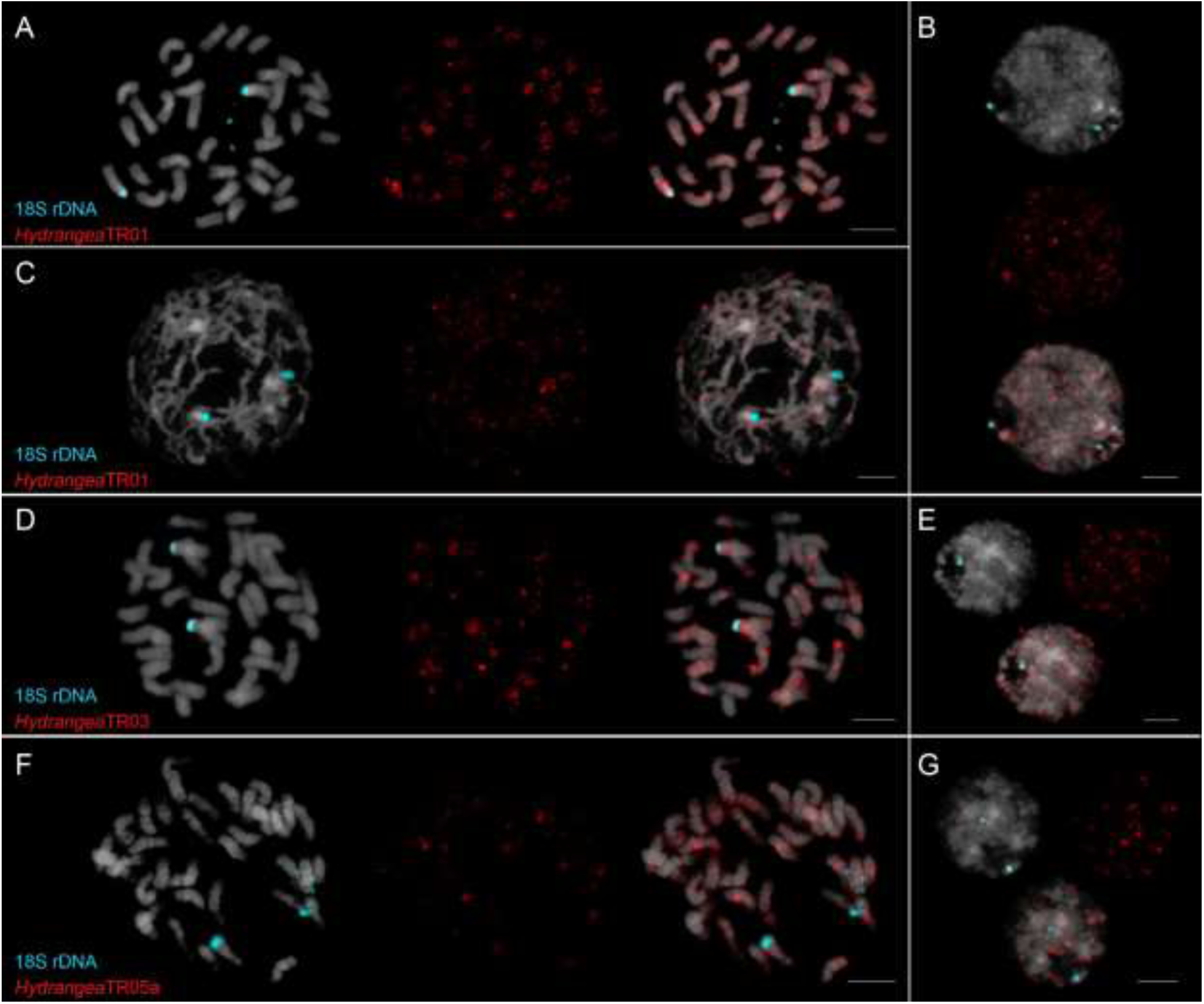
Multicolor fluorescent *in situ* hybridization to chromosome spreads of *H. macrophylla* to determine the chromosomal localization of *Hydrangea*TR01, 03, and 05a. DAPI-stained mitotic chromosomes and interphase nuclei are shown in gray. Turquoise signals indicate the 18S rDNA loci. Metaphase (A, D, F) and prophase (C) nuclei display the dispersed occurrence of HydrangeaTR01, 03, and 05a along all 36 *H. macrophylla* chromosomes. Interphase nuclei of *H. macrophylla* show that HydrangeaTR01, 03, and 05a (B, E, and G; red signals) are located in constitutive heterochromatic regions. Information on probe labeling and detection can be found in the methods section. Scale bars = 5 µm.

The two 18S rDNA signals in each mitotic nucleus indicate one chromosome pair (Fig. 7A, C, D, F; turquoise signals). In Fig. 7A and 7F, one 18S rDNA locus each is still decondensed, as shown by a stretched signal, indicating a despiralized chromatin thread. In comparison, in interphase nuclei, the 18S rDNA signals are very concise, indicating a compact state of the corresponding chromatin (Fig. 7B, E, G; turquoise signals).

The three *Hydrangea*TRs show very similar distribution patterns. All three are found on all 36 *H. macrophylla* chromosomes. Generally, the *Hydrangea*TR signals are rather dispersed along the chromosomes with no clear site preference (both intercalary and distal positions). However, all three *Hydrangea*TRs are enriched on the chromosome arms containing the 18S rDNA. Furthermore, strong *Hydrangea*TR signals were often found near the centromeres, potentially contributing to the (peri-)centromeric regions of these chromosomes (Fig. 7A, D, F). The dispersed distribution pattern of *Hydrangea*TR01 arrays is particularly visible in the prophase nucleus in which *Hydrangea*TR01 signals can be found all along the chromatin threads (Fig. 7B, red signals). As for *Hydrangea*TR01 and 03, most chromosomes harbor several strong signals, indicating major *Hydrangea*TR sites with presumably longer arrays, whereas the minority of chromosomes shows only few, faint signals (Fig. 7A, D; red signals). The opposite applies for *Hydrangea*TR05a: most chromosomes harbor only few, faint signals, indicating shorter *Hydrangea*TR05a arrays, whereas the minority of chromosomes shows several strong signals, indicating longer arrays (Fig. 7F; red signals). Interphase nuclei demonstrate that the *Hydrangea*TRs are preferentially located in the DAPI-positive heterochromatic regions (Fig. 7B, E, G).

## Discussion

To better understand the basis of the genomic variability among the hortensias, we chose a genotype panel of five *Hydrangea* species that covers a wide spectrum of different geographic origins, morphologies, base chromosome numbers, and ploidy levels. This panel serves as a basis to analyze genomic changes over evolutionary timeframes and has the potential to inform hortensia breeding with regards to integration of wild germplasm resources. To develop a robust and useful dataset, we identify (1) plastomes for the evolutionary framework, (2) genome sizes and repeatomes to understand the overall genomic variation, and (3) chromosomal positions of selected repeat families.

### A phylogenetic framework for the 6-hortensia panel

Despite the many efforts to elucidate comprehensive phylogenies within the genus *Hydrangea* (e.g. Samain *et al*., 2010; De Smet *et al*., 2015; Fu *et al*., 2019; Raman *et al*., 2023), this is impeded by the vast diversity of *Hydrangea* species, subspecies and varieties. Therefore, we used plastome data to reconstruct a phylogeny, reflecting the relationship of the precise individuals used in this study. This phylogeny is in line with other current phylogenies (i.e. Raman *et al*., 2023), dividing the analyzed species into two clades, one of them comprising the Japanese diploid species (including the undesignated *H*. spec.) and the other one comprising the polyploid and non-Japanese species. Depending on the fraction of sequence data (e.g. protein coding versus full plastomes) and algorithm, incongruences became visible only in the relation of the three *Macrophyllae* members (*H. macrophylla*, *H. serrata*, and *H*. spec.) to each other, which is due to the high degree of similarity of their plastomes and thus very short branch lengths.

### *Hydrangea* genome sizes vary with chromosomal setup changes and repeatome dynamics

Genome size estimation of two Japanese diploid *Hydrangea* species demonstrated that *H. macrophylla* (2127±273 Mb/1C) has a considerably larger genome than *H. serrata* (1660±108 Mb/1C). This was somewhat unexpected given their close relationship and the fact that both species have the same number of chromosomes (Cerbah *et al*., 2001). Indeed, Zonneveld *et al*. (2005) measured a genome size for *H. serrata* (2102 Mb/1C) that is much more similar to that of *H. macrophylla* than the present estimation indicates. Thus, the question arises where such a difference may come from. It is unlikely that this observation is due to methodological inaccuracies since the genome size estimation for all hortensia individuals analyzed here was carried out with the same flow cytometer under the same conditions and even multiple measurements resulted in the same estimates. Hence, another possible explanation may apply: Genome sizes can vary among individuals of the same species and can be associated with various environmental conditions of the respective regions of origin, such as altitude, temperature, light, humidity, and soil (Bancheva and Greilhuber, 2006; Chalup *et al*., 2014; Inceer *et al*., 2018; Benhizia *et al*., 2021). The growing region of *H. macrophylla* is characterized by milder temperature seasonality in comparison to the more severe temperature changes in the snowy northern areas of Japan where *H. serrata* is growing (Iwatsuki *et al*., 2001). However, the repeat fraction within the genomes may also play a role since we found more repetitive elements within the larger genome of *H. macrophylla* compared to that of *H. serrata*. It may be worthwhile to analyze further *H. serrata* genotypes since the genome size estimates from this study together with those from Zonneveld *et al*. (2005) indicate that there may be a large range in genome size span across *H. serrata* individuals.

The largest genome size among the analyzed species was measured for *H. paniculata* (3484±52 Mb/1C), which is a polyploid species (2n=4x=72 or 2n=6x=108; Funamoto and Tanaka, 1988). Its genome size is about twice as large as that of the diploid *Hydrangea* species, suggesting that the *H. paniculata* variety used in this study may be tetraploid. However, apart from the ploidy, the high repeat content (the highest among the analyzed Hortensias) also increases the genome size of *H. paniculata*.

Most *H. strigosa* accessions have 2*n* = 34 chromosomes, however, there are also a few individuals with 36 chromosomes (Mortreau *et al*., 2010). The relatively small genome size observed for *H. strigosa* (1428 Mb/1C) may therefore be related to the loss of two chromosomes including considerable parts of the genome. This assumption is supported by the fact that for *H. strigosa* accessions having 36 chromosomes, larger genome sizes were estimated in comparison to the aneuploid accessions (Mortreau *et al*., 2010).

The genome size of *H. arborescens* (770 Mb/1C) estimated in this study is the smallest genome size measured for a *H. arborescens* accession so far (Plant DNA C-values Database; 2023/08/17). More comprehensive analyses focusing on the *H. arborescens* intra-species diversity are required to determine whether the reduced genome size of the *H. arborescens* variety “Annabelle” may have arisen during the breeding process from the wild *H. arborescens* ancestor.

### The LTR retrotransposon composition may serve to distinguish *Hydrangea* genomes

In comparison to other plant genomes (e.g. 73% in *Quinoa* and 68% in *Larix*; Heitkam *et al*., 2020, 2021) the analyzed *Hydrangea* genomes show rather moderate overall repeat proportions (42.15% up to 61.07%). However, given that diverged repeats escape the clustering threshold, the actual repeat content is likely to be higher. Repetitive elements are most abundant in the polyploid *H. paniculata* genome, whereas the *H. arborescens* genome is composed of the fewest repetitive elements (Table 2, Fig. 3), highlighting the correlation between repeat content and genome size that was observed for hortensias similar to several other plants (e.g. *Fabeae*, *Solanum*, and *Beta* sp.; Macas *et al*., 2015; Gaiero *et al*., 2019; Schmidt *et al*., 2023).

The repeat composition differs among the six *Hydrangea* representatives, dividing the Japanese diploids (*H. macrophylla* and *H. serrata*) and *H*. spec. from the polyploid *H. paniculata*, Chinese *H. strigosa*, and American *H. arborescens*, which is well in line with the plastome-based phylogeny of these six individuals.

In addition to the six hortensia individuals from this study, the repeatome of the *H. macrophylla* cultivar ‘Sir Joseph Banks’ (GenBank accession number PRJEB32928) was analyzed as well, since this is the first hortensia that was brought to Europe (Tränker *et al*., 2019). We found that its repeat profile is almost identical to that from the Japanese cultivar *H. macrophylla* f. *normalis* used in this study (see Supplementary Data Fig. S4). Therefore, and due to the fact that the *H. macrophylla* cultivar ‘Sir Joseph Banks’ is considered to be the ancestor of the entire European hortensia breeding (Tränker *et al*., 2019), following conclusions drawn based on the *H. macrophylla* f. *normalis* repeatome may also apply to most European *H. macrophylla* cultivars. The undesignated *H*. spec. also shares nearly all repeat clusters with the two Japanese diploid *Hydrangea* species (*H. macrophylla* and *H. serrata*), including a similar respective repeat abundance. In combination with the plastome-based phylogeny that groups *H*. spec. closely with *H. macrophylla* and *H. serrata* (Supplementary Data Fig. S2), this indicates that *H*. spec. is a member of the section *Macrophyllae*. This suggests that the repeat profiles may be employed to ascertain the conspecificity of undesignated varieties. Interestingly, two repeats specific to *H*. spec. were observed, indicating minor differences in the repeat profiles among the *H. macrophylla* representatives that may be used to distinguish even closely related species or varieties (Fig. 5). Further analysis of these specific repeats may help to infer phylogenetic trajectories and to trace hybridization events (during cultivation). In other words, further repeatome analysis may assist to classify unknown varieties within these species.

As in most plant genomes (reviewed by Weiss-Schneeweiss *et al*., 2015; Orozco-Arias *et al*., 2019), LTR retrotransposons are the most abundant repeats in *Hydrangea* genomes, accounting for 31-51%. Members of the Ty3/Gypsy superfamily, especially Tekay elements, are the predominant repeats among them. In terms of genome share, the polyploid *H. paniculata* genome contains approximately four times more Tekay elements (963 Mb/1C) than the diploid Japanese species (218-282 Mb/1C) (Fig. 4, right). This increased abundance may indicate the acquisition of new Tekay elements as well as an amplificational burst of already present ones after the polyploidisation event as demonstrated by the comparative repeatome analysis.

In contrast, Ogre and Athila elements are less abundant in the polyploid *H. paniculata* genome compared to the Asian diploid species (*H. macrophylla*, *H. serrata*, and *H. strigosa*). A possible explanation for this could be the efficient elimination of these elements from the *H. paniculata* genome or an increased Ogre/Athila amplification within as well as the acquisition of new Ogre/Athila elements into the Asian diploid *Hydrangea* genomes. Indeed, we detected a number of clusters annotated as Ogre and Athila retrotransposons, respectively, that specifically appear in the diploid *Hydrangea* genomes, indicating that integration events of Ogre and Athila sequences occurred several times after the split of the Asian *Hydrangea* species from their common ancestor.

Concluding, based on the pattern of shared repeat clusters among the analyzed *Hydrangea* representatives as well as the assumption of a parsimonious evolution, the following scenario may be conceivable: The genome of the *Hydrangea* ancestor comprised a limited set of repeats. The geographic separation of the American *H. arborescens* from the remaining *Hydrangea* species led to the acquisition of distinct repeats including some Tekay and Athila elements. The ancestor of the Asian branch acquired distinct repeats as well, including some Ogre and Retand elements. The subsequent split of the Chinese *H. strigosa* was followed by the acquisition of further Ogre and Athila elements. Within the Japanese branch, a polyploidisation event resulted in the split of the polyploid *H. paniculata* from the remaining diploids. And whereas *H. paniculata* only acquired a few new repeats, the ancestor of the remaining diploids acquired a whole range of new Ogre and Athila elements (among others). Following this premise, the detection of Retand elements may be used to distinguish Asian from American *Hydrangea* species. A separation of Asian *Hydrangea* species based on Ty3/Gypsy retrotransposons may be possible as well. However, the specific retrotransposon families should be further characterized for this purpose.

The LTR retrotransposon composition of the Ty1/Copia superfamily shows similarly high interspecific variability among the *Hydrangea* genomes (Fig. 4, left): Whereas elements of the TORK lineage are the predominant Ty1/Copia representatives within the Asian diploid species (Angela elements within the Japanese diploid *Hydrangea* genomes; TAR elements within the Chinese *H. strigosa* genome), elements of the Retrofit lineage (Ale elements) are most abundant in the Japanese polyploid and the American species, indicating a high retrotransposon turnover during *Hydrangea* evolution. Such a fast evolution is typical for repetitive elements (Ugarkovic *et al*., 2005; Kuo *et al*., 2021). Species-specific Ty1/Copia retrotransposons were also identified (e.g. Oryco/Ivana elements within the *H. strigosa* genome), pointing to the integration of these elements into the *Hydrangea* genomes after *Hydrangea* speciation. Such interspecific differences in the LTR retrotransposon composition have been reported in many flowering plant species, including differences on an intraspecific level (Du *et al*., 2010; Usai *et al*., 2017). However, no species-specific repeats were identified in the *H. macrophylla* genome, suggesting that this genome comprises the fundamental repeat set of Japanese diploid *Hydrangea* species, potentially resembling their common ancestor.

### Potential applications in breeding

*Hydrangea* is an ornamental plant with over 250 years of cultivation history. During the Edo period, the German physician and naturalist Philipp Franz von Siebold had extensively studied and documented various species in his work, ‘Flora Japonica,’ which was published between 1835 and 1870. Even so the genetic studies have been constrained by limited resources. Simple sequence repeat (SSR) markers and single nucleotide polymorphisms (SNPs) have been effectively used in the study of *Hydrangea* species to assess their genetic diversity and relationships. The application of these markers in *H. macrophylla* has highlighted their utility in distinguishing closely related clones. Genetic diversity in 114 taxa of *H. macrophylla* were analyzed using SSR markers (Reed and Rinehart. 2007). Wu *et al*. (2021) conducted a transcriptomic study on the cultivars to discover SSR markers in *H. macrophylla*. Nashima *et al*. (2021) identified specific genetic markers using SNPs linked to the double flower loci of *H. macrophylla*. These studies utilized genome sequence data to identify DNA markers, demonstrating their valuable roles in advancing the knowledge of *Hydrangea* genetics, which is crucial for breeding programs and cultivar identification. Our repeatome analysis would provide valuable insights by identifying specific genetic markers, such as SNPs from highly variable regions of repeat families like retrotransposable elements, through low-coverage NGS sequencing. Reiche *et al*. (2021) developed molecular markers for the identification and differentiation of poplar genotypes using Inter-SINE Amplified Polymorphism (ISAP). These markers can help pinpoint potential parents and ancestors and important genes for specific traits. Plant breeders can use this genetic information to analyze inheritance patterns in progeny, aiding in breeding, conservation, and improvement of this ornamental species.

### The two *Hydrangea* clades show differing satDNA profiles

Conserved and specific satDNAs among the six hortensias were identified using the comparative clustering analysis. The identification of satDNAs is a fundamental step to compare the genetic diversity in plant species since satDNAs evolve fast and are therefore suitable as cytogenetic markers (Ugarkovic *et al*., 2005; Kuo *et al*., 2021).

The Chinese *H. strigosa* and American *H. arborescens* genomes are characterized by a lower satDNA variety and overall lower satDNA abundances compared to the Japanese *Hydrangea* genomes. Similar to the retrotransposon composition, the analyzed *Macrophyllae* members (*H. macrophylla*, *H. serrata*, and *H*. spec.) share most of their satDNAs. The appearance of genome-specific satDNAs (e.g. *Hydrangea*TR02 and 10 in the *H. serrata* genome) indicates a fast satDNA evolution, especially since the split of the Japanese diploid *Hydrangea* species (including *H. serrata*) occurred rather recently (within the range of 0.58-2.53 million years ago; Raman *et al*., 2023). Similarly, the species-specific satDNA amplification and the emergence of new monomer variants (e.g. *Hydrangea*TR05b) are hallmarks of the evolutionary drive.

The fact that the *H. serrata*-specific satDNAs *Hydrangea*TR02 and 10 were not detected in the genome of the undesignated *H*. spec. give rise to the assumption that *H*. spec. represents another *H. macrophylla* variety. Thus, the *H. serrata*-specific satDNAs *Hydrangea*TR02 and 10 may help to assign undesignated *Macrophyllae* species, even though the members of this section are genetically very close (Raman *et al*., 2023) and cannot even be resolved using an otherwise very robust plastome-based phylogeny (Supplementary Data Fig. S2).

### Major tandem repeats in *Hydrangea* are distributed throughout the chromosomes

A major 18S rDNA locus was detected on two *H. macrophylla* chromosomes forming the secondary constriction site (Fig. 7). This observation is consistent with previous studies (Van Laere *et al*., 2008). The three most abundant tandem repeats in the *H. macrophylla* genome, *Hydrangea* TR01, 03, and 05a, were dispersed throughout the *H. macrophylla* chromosomes. Their distribution patterns are very similar. *Hydrangea*TR01 and 03 showed a high *Hydrangea*TR abundance on most chromosomes, whereas most *Hydrangea*TR05a chromosomes harbor only few, marginal signals, indicating a low *Hydrangea*TR abundance. These signal intensities corroborate the bioinformatics results regarding the respective satDNA abundances.

Highly repeated distribution patterns at specific chromosome regions suggest that *Hydrangea* TRs may be involved in the maintenance of the heterochromatic structure, which is commonly observed in eukaryotic genomes including *A. thaliana* and rice (Garrido-Ramos, 2015). Large arrays of tandemly repeated sequences found in (peri)centromeric regions have also been detected in a wide range of species (Heslop-Harrison and Schwarzacher, 2011). All three TR signals were also co-localized with rDNA signals, suggesting that *Hydrangea*TRs are also interspersed in the rDNA array.

The large and small localization on all chromosomes suggested that *Hydrangea*TRs in the *H. macrophylla* genome may frequently change their structures during plant evolution. Transitions from transposable elements to satDNAs have been observed in some plant species, such as satDNAs in maize and *Solanum bulbocastanum* derived from LTR retrotransposons (Meštrović *et al*., 2015). SatDNAs and transposable elements are present in gene-poor heterochromatic regions, where the repeat copy number variation should not affect the genome stability. Therefore, *Hydrangea*TRs may have been generated as a result of their location adjacent to transposable elements in the *H. macrophylla* genome and may have accumulated at each chromosomal site by subsequent amplification.

Using bioinformatics (*i.e*. the RepeatExplorer2 pipeline) the three *Hydrangea*TRs were found in other *Hydrangea* species as well. The comparison with other Japanese diploid species may result in different signal patterns during FISH, *i.e*. if there are aberrations in the number of chromosomes. Other *Hydrangea*TRs which were found to be species-specific have the potential to be used as chromosome markers as well. Chromosome-specific satDNAs were identified and used for karyotyping the 34 chromosomes of *Crocus sativus* (Schmidt *et al*. 2019). Paesold *et al*. (2012) identified chromosome-specific markers for discriminating all nine chromosomes of *Beta vulgaris*, which were associated with centromeric, intercalary, and subtelomeric regions of the chromosomes. Mortreau *et al*. (2010) reported the karyotyping for *H. involucrata* (2n=30), and *H. strigosa* (2n=34) by FISH using 5S and 18S-5.8S-26SrDNA and chromomycin A3 fluorochrome banding. The species/group-specificity and repeat abundance of *Hydrangea*TRs 02, 04, 10 and 13 make them promising candidates to serve as markers for further cytogenetic and phylogenetic analyses.

## Conclusions

Repeatome studies are essential for understanding plant genetic diversity. We performed a comparative repeatome analysis in *Hydrangea* using modern bioinformatic techniques and FISH-based chromosome mapping. Investigation of plastid genome polymorphism revealed genetic differences among *Hydrangea* lineages. The composition of tandem repeats suggested a closer relationship among Japanese diploid species. Further repeatome analysis may assist in classifying unknown *Hydrangea* varieties. The development of cytogenetic and molecular markers based on repeatome and plastome analyses is crucial for breeding and organizing genetic resources. Identifying specific sequences in Asian *H. macrophylla*, *H. serrata* and American *H. arborescens* can help manage world genetic resources. This comprehensive genomics analysis, including nuclear genome repeats, plastid and chromosome dynamics, is essential for future studies of Hydrangea genetic diversity.

## Acknowledgements

This work was supported by the networking and exchange grant “Accessing complementary hortensia germplasms to enable floricultural genomics” by the DAAD via funding from BMBF and the JSPS to NO, SW and TH (DAAD 57607355 and JPJSBP-120223504). Computational resources for RepeatExplorer2 analyses were provided by the ELIXIR-CZ project (LM2015047), part of the international ELIXIR infrastructure. Furthermore, this work was supported by a postdoc fellowship of the German Academic Exchange Service (DAAD) granted to MJ. To other colleagues, Ms. Hina Shimomai (Kobe Univ) and Luise Keßler (Technische Universität Dresden), we would like to express our gratitude for experimental support and fruitful discussion.

## Notes

### Competing Interest Statement

The authors have declared no competing interest.

