## Supplemental data for "Repeatome landscapes and cytogenetics of hortensias provide a framework to trace *Hydrangea* evolution and domestication"

* equally first

+ equally corresponding

1 Graduate School of Human Development and Environment, Kobe University, Nada-ku, Kobe, 657-8501, Japan

2 Faculty of Biology, Technische Universität Dresden, D-01069 Dresden, Germany

3 Institut für Ökologie, Evolution und Diversität, Goethe-Universität Frankfurt, 60438 Frankfurt am Main, Germany

4 Departamento de Botánica, Instituto de Biología, Universidad Nacional Autónoma de México, Mexico City, Mexico

5 Abteilung Botanik und Molekulare Evolutionsforschung, Senckenberg Gesellschaft für Naturforschung, 60325 Frankfurt am Main, Germany

6 Institute of Biology, NAWI Graz, Karl-Franzens-Universität, A-8010 Graz, Austria

### Supplementary Data

### Supplementary Data

#### Supplementary Data Figure S1

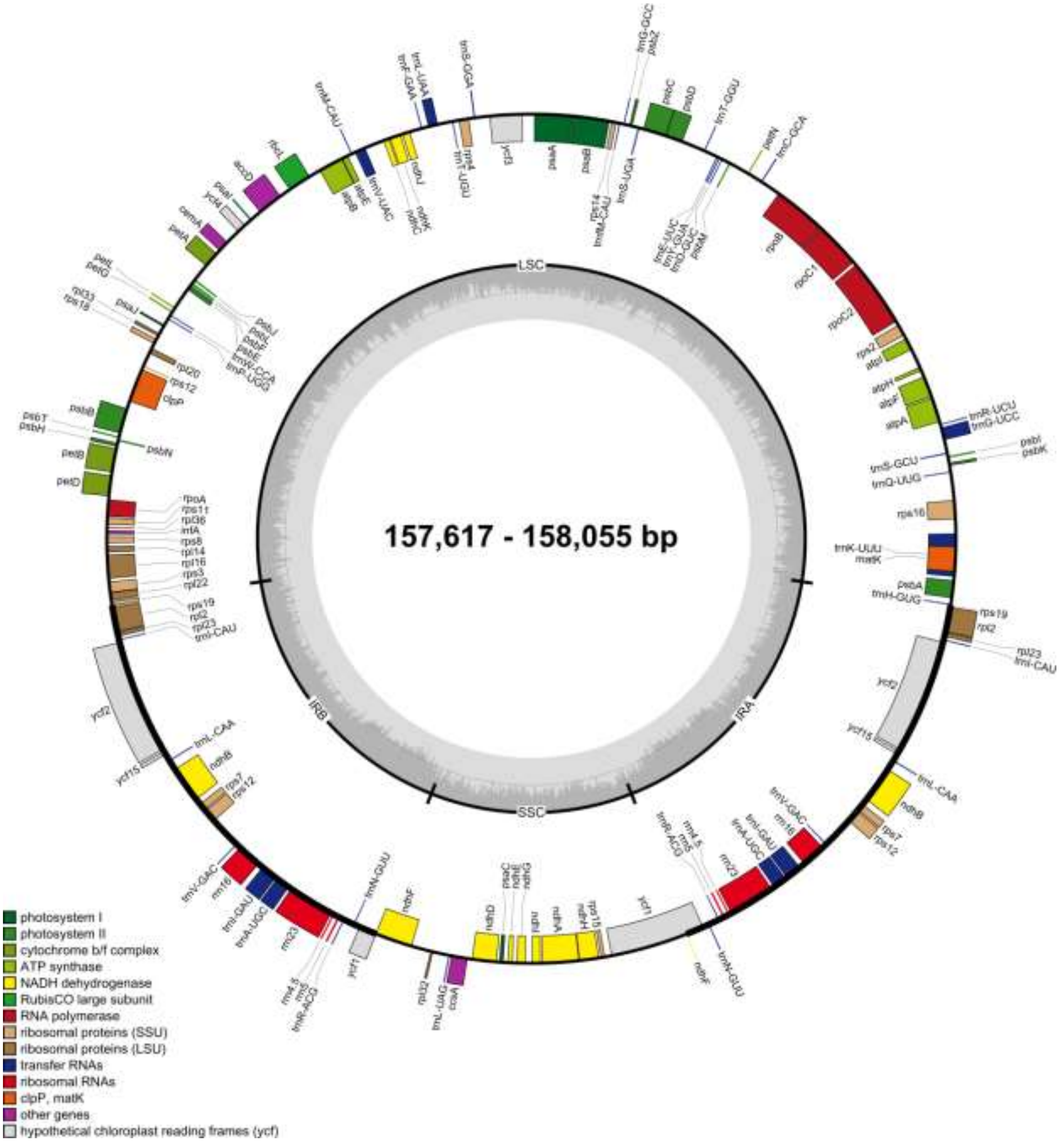

### Supplementary Data

**Fig. S1: Circular plastid genome map of the six *Hydrangea* genotypes used in this study.** Different colors indicate genes of different functions. The genes inside the outer circle are transcribed clockwise and the genes on the outside are transcribed counter clockwise. The large single copy (LSC) region, small single copy (SSC) region, and two inverted repeat (IRA and IRB) copies are indicated on the inner circle. The gray circle visualized the GC content across the different plastome regions.

### Supplementary Data

#### Supplementary Data Figure S2

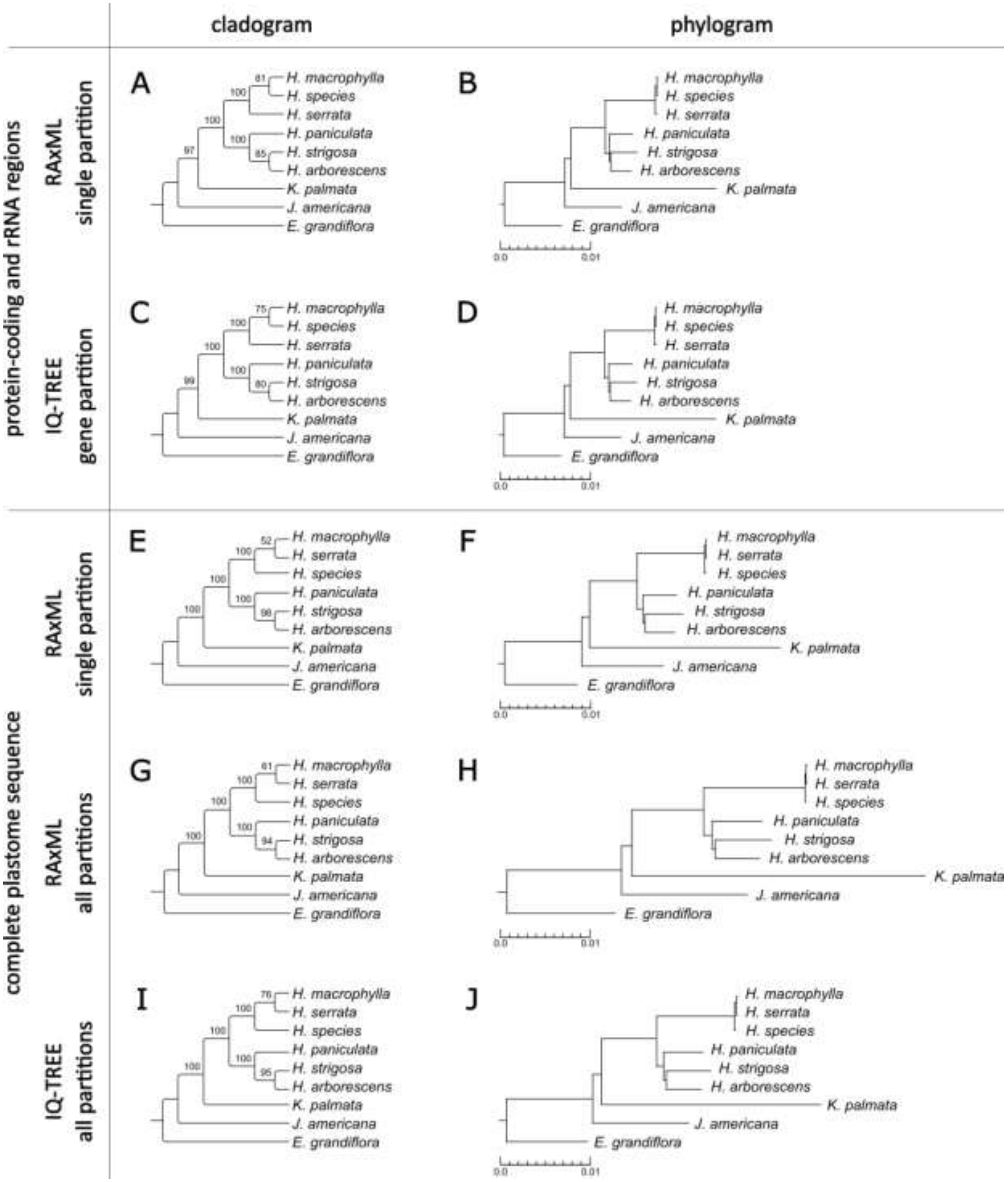


**Fig. S2: Plastome-based tree reconstructions of the studied *Hydrangea* genotypes using both RAxML (A, B, E, F, G, H) and IQ-TREE (C, D, I, J).** The topologies were calculated based on the protein-coding and rRNA regions (single partition: A, B; gene partition: C, D), as well as on the complete plastome sequence excluding one IR copy (single partition: E, F; all partitions [genes, introns, spacer]: G-J). On the left side (A, C, E, G, I), the cladograms highlighting the bootstrap values are depicted. Support values are based on 1,000 bootstrap replicates. On the right side (B, D, F, H, J), the corresponding phylograms, drawn to scale, are shown. The outgroup sampling is comprised of representatives from the sister tribe Philadelpheae (*Kirengeshoma palmata*: NC_044808) and the sister subfamily Jamesioideae (*Jamesia americana*: NC_044836). *Eucnide grandiflora* (NC_044767), a member of sister family Loasaceae, was used for rooting.

#### Supplementary Data Figure S3

### Supplementary Data

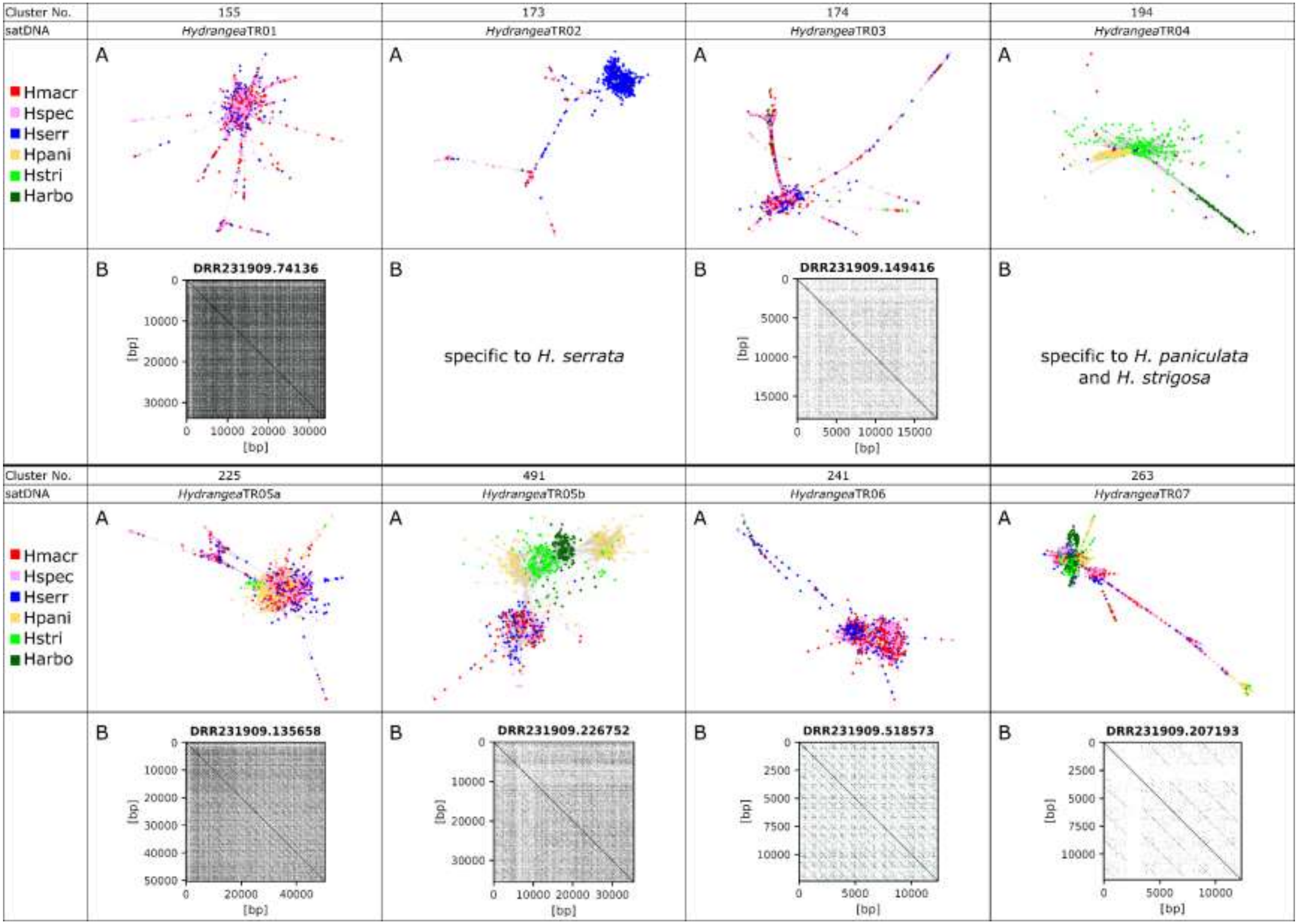

### Supplementary Data

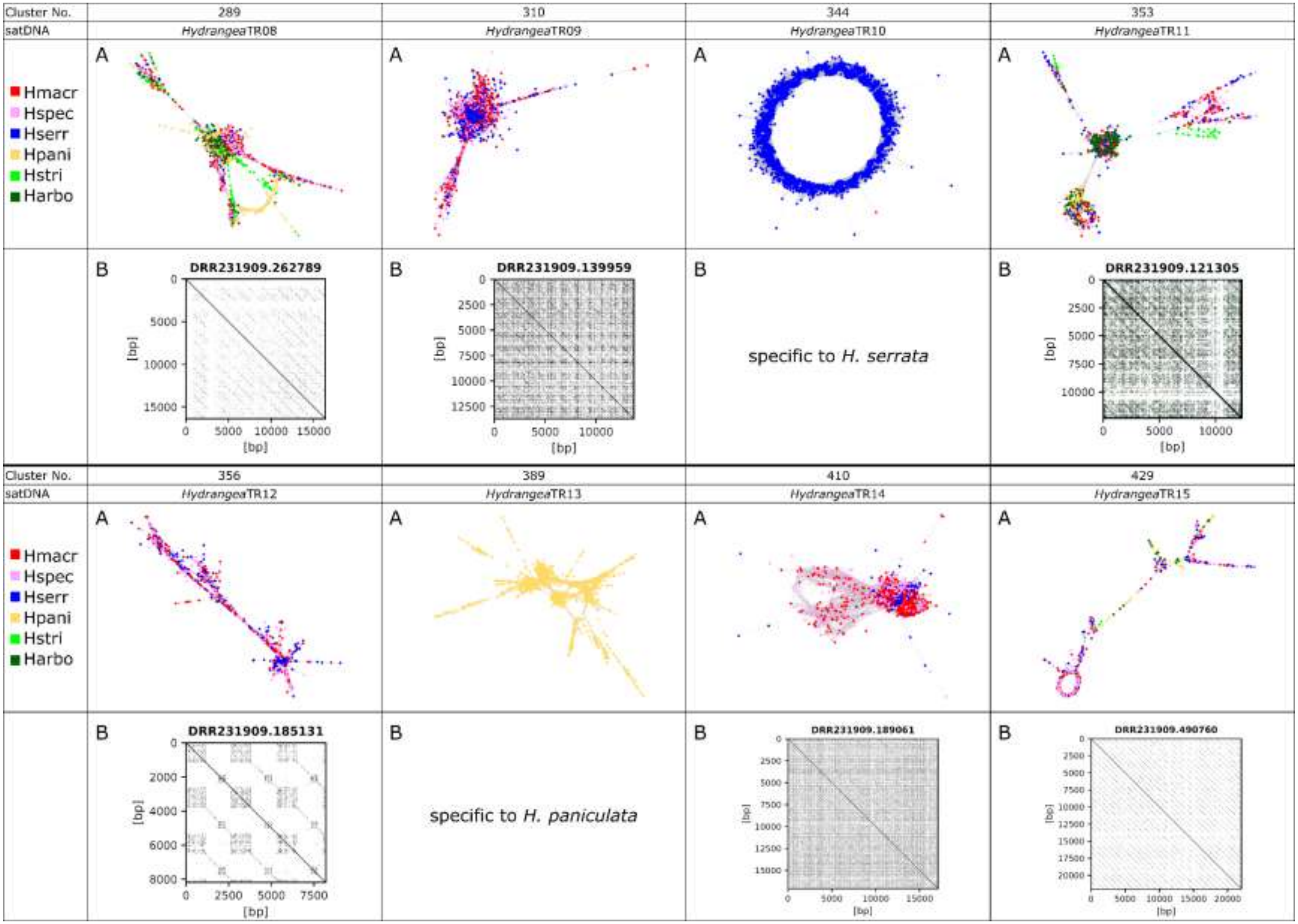

### Supplementary Data

**Fig. S3: *Hydrangea*TR characteristics determined by RepeatExplorer2 and self-dotplot analyses.** (A) SatDNA-typical bulk or ring-shaped cluster graphs resulting from the comparative RE2 analysis of the six different *Hydrangea* genotypes. The origin of the distinct reads is color-coded and highlights *Hydrangea*TR specificities. (B) Arrangement of *Hydrangea*TR monomers on long reads of *H. macrophylla* (DRX222164). *Hydrangea*TRs that are specific to other *Hydrangea* species could not be detected in the *H. macrophylla* dataset. Forward matches are marked by black lines, whereas reverse matches are indicated in green. To account for the ONT read error rate of 5-10%, we chose a word size of 20 bp and allowed five mismatches.

#### Supplementary Data Figure S4

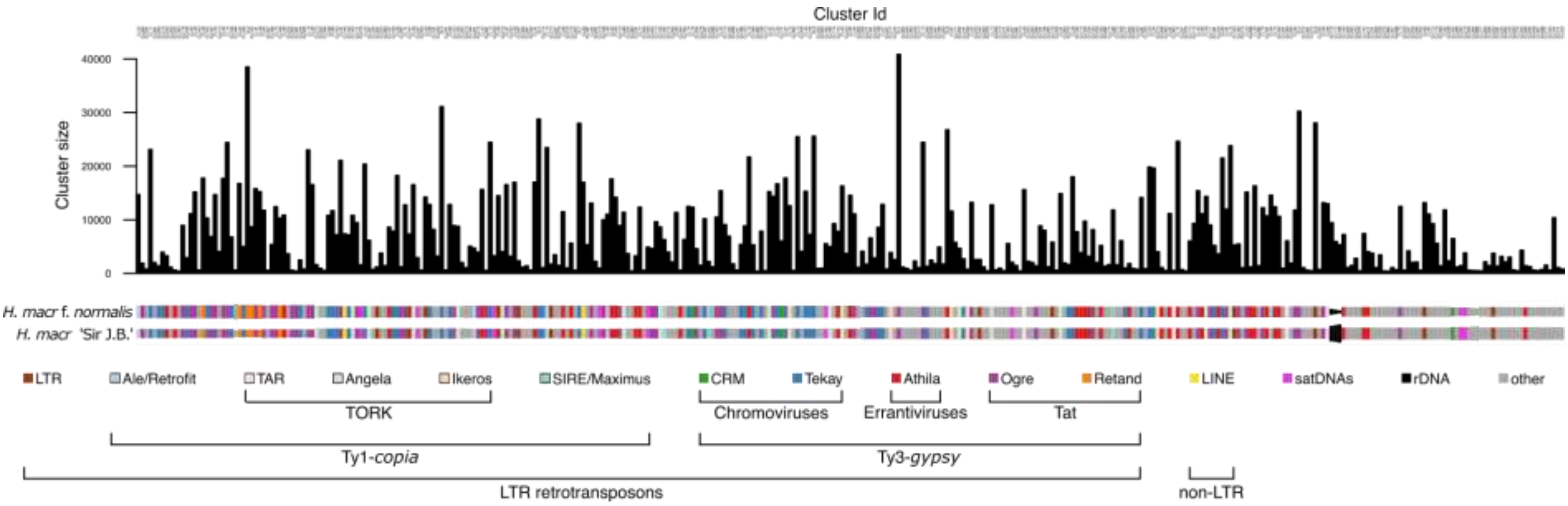


**Fig. S4: Comparative repeat composition between two *H. macrophylla* cultivars, *H. macrophylla* f. *normalis* (used in this study) and *H. macrophylla* 'Sir Josef Banks' (main basis for all *Hydrangea* breeding in Europe; Tränker *et al.*, 2019).** The bars represent the distribution of clusters comprising at least 432 reads ( ≥ 0.01% of the analyzed reads) among the analyzed species. Rectangles are colored according to the type of repetitive element and their size is proportional to the genomic abundance in the respective cultivar.

### Supplementary Data

#### Supplementary Data Table S1

**Table S1: Primer sequences for the amplification of different *Hydrangea*TRs.** *Hydrangea*TR01, 03, and 05a were amplified using genomic DNA of *H. macrophylla*.

| Primer name | FW Sequence | RV Sequence | Ann. Temp. [°C] |
|---|---|---|---|
| *Hydrangea*TR01 | CGT TAT ATT CTC AAA ACG AGG G | CGG TTC TCG CAA CAG ATT TG | 55 |
| *Hydrangea*TR03 | CCG ATT CAC TAC TCA AAA CCT CA | AAC GCC ACG AAA CCT TAC AC | 55 |
| *Hydrangea*TR05a | TCA AAA TGC AAA CCG TCG AT | GAG TCC GAT TCC CGT TCC | 53 |

#### Supplementary Data Table S2

**Table S2: Read datasets used for repeatome analyses and the reconstruction of plastome sequences.** The reads were sequenced using the Illumina Novaseq 6000 system as 150 bp paired-end reads for each genotype and published as part of the study ERP151402.

| Genotype | Accession no./ENA Run ID (Illumina reads) | Experiment (Illumina reads) | Accession no. of the corresponding plastome sequence |
|---|---|---|---|
| *H. macrophylla* | ERR12526782 | ERX11900996 | OR701877 |
| *H.* spec. | ERR12526783 | ERX11900997 | OR701880 |
| *H. serrata* | ERR12526784 | ERX11900998 | OR701879 |
| *H. paniculata* | ERR12526785 | ERX11900999 | OR701878 |
| *H. strigosa* | ERR12526786 | ERX11901000 | OR701881 |
| *H. arborescens* | ERR12526787 | ERX11901001 | OR701882 |

#### Supplementary Data Table S3

**Table S2: Absolute amount [Mbp] of different repeat classes in six *Hydrangea* genotypes.** Abbreviations as in Table 1.

| Repeat | | Lineage | Clade | Genome Fraction [Mbp] | | | | | |
|---|---|---|---|---|---|---|---|---|---|
| | | | | **Hmacr** | **Hserr** | **Hpani** | **Hstri** | **Harbo** | **Hspec** |
| **LTR retro-trans-posons** | **Ty1/Copia** | Retrofit/Ale | | 30.63 | 26.29 | 84.66 | 17.65 | 8.09 | 31.63 |
| | | Alesia | | 0.00 | 0.00 | 0.00 | 0.15 | 0.00 | 0.00 |

#### Supplementary Data

| | | | | | | | | | |
|---|---|---|---|---|---|---|---|---|---|
| | | Oryco/Ikeros | | 15.74 | 11.02 | 16.37 | 7.97 | 2.70 | 14.58 |
| | | TORK/Tork | | 2.13 | 1.99 | 23.34 | 8.18 | 1.08 | 1.33 |
| | | TORK/Angela | | 92.74 | 63.82 | 83.27 | 7.02 | 8.86 | 70.46 |
| | | TORK/TAR | | 15.53 | 8.92 | 20.90 | 16.97 | 2.54 | 14.58 |
| | | TORK/Ivana | | 0.00 | 0.30 | 1.05 | 2.38 | 0.00 | 0.00 |
| | | SIRE | | 13.61 | 6.89 | 12.19 | 5.15 | 0.85 | 11.55 |
| | | Bianca | | 1.49 | 1.57 | 8.01 | 5.17 | 0.46 | 1.33 |
| | | **Total Ty1/Copia** | | **171.86** | **120.81** | **249.80** | **70.65** | **24.56** | **145.46** |
| | **Ty3/Gypsy** | chromovirus | CRM | 12.76 | 11.13 | 17.42 | 7.58 | 2.93 | 11.55 |
| | | | Tekay | 282.68 | 218.18 | 963.33 | 251.52 | 148.23 | 223.87 |
| | | Non-chromovirus | Athila | 183.56 | 130.24 | 105.22 | 25.69 | 45.58 | 129.74 |
| | | | Ogre | 295.23 | 213.17 | 169.67 | 233.64 | 4.08 | 249.82 |
| | | | Retand | 65.30 | 56.20 | 101.04 | 45.16 | 2.16 | 58.71 |
| | | **Total Ty3/Gypsy** | | **839.53** | **628.92** | **1356.67** | **563.60** | **202.97** | **673.70** |
| | Unclass. LTR | | | 83.17 | 47.97 | 90.24 | 41.27 | 9.78 | 17.27 |
| | **Total LTR** | | | **1094.55** | **797.70** | **1696.71** | **675.52** | **237.31** | **836.39** |
| LINE | | | | 11.06 | 7.80 | 24.39 | 3.43 | 4.85 | 8.33 |
| Pararetrovirus | | | | 2.98 | 0.00 | 6.62 | 1.14 | 0.00 | 0.00 |
| | **Total retrotransposons** | | | **1108.59** | **805.51** | **1727.72** | **680.09** | **242.17** | **844.72** |

### Supplementary Data

| | | | | | | | | |
|---|---|---|---|---|---|---|---|---|
| **DNA transposon** | TIR | EnSpm_CACTA | 0.43 | 0.00 | 3.83 | 0.00 | 0.08 | 0.38 |
| | | hAT | 1.91 | 0.67 | 8.71 | 0.56 | 0.15 | 2.84 |
| | | MuDR_Mutator | 2.77 | 0.94 | 9.06 | 1.81 | 0.31 | 2.27 |
| | | PIF_Harbinger | 0.00 | 0.00 | 3.14 | 0.00 | 0.00 | 0.00 |
| | Helitron | | 0.00 | 0.00 | 0.70 | 0.00 | 0.00 | 0.00 |
| | **Total DNA transposons** | | **5.10** | **1.60** | **25.43** | **2.37** | **0.54** | **5.49** |
| **Tandem repeats** | satDNAs | | 24.67 | 39.39 | 19.86 | 15.97 | 1.85 | 24.81 |
| | rDNA | 35S | 4.89 | 16.37 | 47.03 | 9.71 | 7.01 | 7.77 |
| | | 5S | 0.00 | 1.59 | 0.35 | 1.53 | 0.23 | 0.00 |
| Unclass. repeats | | | 134.00 | 126.46 | 307.29 | 133.12 | 72.69 | 168.76 |
| **Total repeats** | | | **1276.84** | **990.85** | **2128.03** | **842.66** | **324.56** | **1051.55** |

### Supplementary Data

#### Supplementary Data S1:

**Data S1: *Hydrangea*TR nucleotide sequences derived as RepeatExplorer2 consensuses from the comparative analysis.** The start and the end of the monomer is arbitrary. Primer binding sites (corresponding to the primer sequences listed in Table S1) are accentuated by bold, underlined fonts.

>HydrangeaTR01-consensusmonomer_60bp
C**CGTTATATTCTCAAAACGAGGG**TTAACG**CAAATCTGTTGCGAGAACCG**AGAATATAAAT

>HydrangeaTR02-consensusmonomer_790bp
AAATATAGGTACAACATGCCATAGCATAGGAACAGTATGGTATAGCCAAGATATTGGATGAAAA
CAATACTCTTAAAGACTATTCCTGGAGATAGTCTCTTTGTCTACGTACACCAGGTTCCCGAATA
CAGACTCTTAAAGCAAATTCCAGGAGATAGTCTCTGATTATATTTGATTATCCATTCCAGAGAC
AAACTATGTCATTTTCTATTTTACATATTGGATACAAAGATACCTATTGTAAGACTTTTTTCAT
AGACAGTCTAGTTATGTGTCTTTGGCCAGACTGTATTTCTGTTAGCATATCTGCATTGTCCAGC
GATAAAAAAAATACACAAACAGATAAACAACAAAATCACATACCTCTTATACCAAAATGATAAC
ATATCAAAGAACTGAAAGTTGTAAGACTTTTGAGATGAGGACTTTGAGATTTGAAATTCAAAGT
TCAGTAGTGAGAAACTTAAATTTTTGGACAAAACAGTTTTTGGTTCTTTTTAACATGATGATTT
ATGGGATGAAAATAGCACTTCCAAAGACTATTCCCGGAGATAGTCTCAATAAAACTCTTTCGAA
AAACTATTCGCGGTGATATTCTCTTAGTTTTTGTACAACATGGCATAGTTTACATACATGATGA
TACATCAAATAATAGTATAAGTACAATGTGGTACAACCTTGTACTGCATGGCATGTTATACCGT
AGTAGAGTGTAAGTATAACATGCTATGTCATATTATAGTGTAGGTATAGTGTGTTATAGTATAG
GTTCAATATTTTTTTCAAAAAA

>HydrangeaTR03-consensusmonomer_179bp
TTA**CCGATTCACTACTCAAAACCTCA**AAAACTAAGCTTTTTCAAGCCTAAGTGACTAAAGAGTC
ATTTTTTCGCATTTTTTGTGAACCGTGAATCCGTTATGCATTCCGTTAGTTCCCGCAAGTTTAT
TTAGGCATGTTTGAGGTGC**GTGTAAGGTTTCGTGGCGTT**TGGTTAAGCGGT

>HydrangeaTR04-consensusmonomer_172bp
GTGTAAGTGCAAAACATAAATGTCGAGTTTTCACCCGACCCTCACACACAAGTTATAAAACACC
AAAATACACGTGTTCAAATTGACTTTCTAGAAAAACTTGACTGGTGAGAGGTAGTTCGTGAGTT
ATTTGACCCGAAATGGCCTTTTCGGCAATTTTTAAAATTGGAGG

>HydrangeaTR05a-consensusmonomer_178bp
CACTACAAAACAACTTTTTTACCTCGTAGACTATGAAAAACTAAGTTTTTGGGGTTTGGGGGG**T**
**CAAAATGCAAACCGTCGAT**CCGGTAACCACGAAACTTAACTCGCACCTCCCACACGTCTAAAGA
AAGTTGTAGGAAGAAAC**GGAACGGGAATCGGACTC**CCGGTTATCAAGATA

>HydrangeaTR05b-consensusmonomer_178bp
CGCGAAAAAATAGAATTTCAAACACTTAGGCTTTAGAAAACTTAGTTTTGGGGTTTTGACGGGT
GAAACGGTGAACCGCCAATCGGAATGACACGAAACTTGACCGACACCTCACACACGCCTAAAGA
AAGTTGTAGGAACAAACGGGACCCTTATCGGATTCCCGGTTTGAGAGTTA

>HydrangeaTR06-consensusmonomer_157bp

### Supplementary Data

GGTATAATTAACAAATATGGGGAATAATGAACAAATATGCAACATGGGAGATGACAACTATTGT
AGGGGAATCCCCCACAATTGTAAAGAATCCCCCACAGTTTGTCTAGAGTCCCCCACAAACTGTT
GTATATGGGGGATTCTTAACAAACTGTGG

>HydrangeaTR07-consensusmonomer_179bp
TCTCTCAAAACCTCAAAACTTAGTTTTAGAAGGCCTAAGTCTATGAATTTGCGTATTTGCGTCT
AACTTGCAATCCGTGAGTCGGAATTGCGTGCCGTTTCTTCCTACAAGTCTATTTAGGTGAGTAT
GAGGTGTAGGTTCCAGTCTCGTGTCATTCCGAGTGCCGGAACCCATTTTGG

>HydrangeaTR08-consensusmonomer_276bp
TGGGGCTTACATTTAATAAGTTAATATAATTAAAGTCTTCAAATAATACGGAGTGACCTACCCC
AAGGTTTTTCGACGTTCTGGATCCAATGGTAGGGTCCATTTGGCCTGAATATATTTGTAAAATG
ACTATTTTGCCCTTGAAACTTTAAAATGCCATAACTTGTTAAGATTAAACCAGAATTGAACGTC
GTCAGAGCCTGTGAGCTCATATTAATATGAAGAACATTTTGAGACCTAAAAAGTACATATATCG
TAAAATGACCAAATTGCCCT

>HydrangeaTR09-consensusmonomer_49bp
TATCGTGAGCTTCCTCACTCCTTTGTCCAAAATCGTGAGCTTTCTCACT

>HydrangeaTR10-consensusmonomer_733bp
TTTGATACTCCATTGACTGTTAGTTGAGTACTATGGGGGGAACACGCCAATCGTTTTTGGACCC
TTGTCACCTCTCCATCTATGTGCACCAACTAATTCTGGAATATAAATAATAAATATACACCCGC
CCCCCAGTCACATACAGTCAAAAATGCTAAATGATTATTCATCTGATTACTCAACAGCCCTCAG
ATCTCTCTATAACTTGAGAGTGTTTTACCTAAAAAAAAGTCACATACAGAGAGTGATTTTTGAG
GGGTTTCTCTAACATAAAATATGCCTCCACAATCCTCAGGTATTGTTCTTGTTCTTCCAGTGAT
CTATTTCTTCGATCTTCTTCTTGCTAATTTTTTGTTCCCCTTTTCTCATCTTTGATATATAAAA
TACTACCCAAATAACCAAATAGATACCCAAATTTAATTACTAGAGTTGCTCTAAGGAAGCATTT
AATGCTTGGCGTGAGGGAGTTGGCAAAAGATATCACGTGAAAGGTGGTTTGACAATTTGGGGCG
CTAGAAACATGCTGAGTACTGGCTACTCTCATTAATGCTGTCACCCCTTTCTTTTGGGAAAAGA
TATCACGTGATTCCCCTATAATCTGTGTTTTTATAATTCCAGAATTTAAATGACCGCGTTTGGG
AAGATGTGGGCAATAATGGGTGCGTGGACTATAGTTCTCCATCAACAAACACATTGGCTAATTT
GGCTGGCTTTCTAGCCAAAAAAAAAATAAA

>HydrangeaTR11-consensusmonomer_316bp
ACGACCCTGGTGTTGTGGGCGGGCTGCTGTAGGTGCCTTTGTGGCTGCAGTAAGTGCTGGCAGG
CTACACACTATGGGAAGTTCGTCCCACATCGCCTGGGTGTGGAAAAGTAATGTGCTATATATGT
GTGGTTCCAAGTCTCCCTAGTAAGAGGCCTTTTGGGTAGTGGCCCAAGAACAAATCCGTGCGGG
TTTGGGCCCAAAGCGGACAATATCTTACTGAGCGAGACTTGGGTCGTGACAGAATGGTATCAGA
GCTAGACCCAGCCGGAAGTGTGCCAACGCGGACGTTGGGCCCTAAGGGGGGTGGAATGTA

>HydrangeaTR12-consensusmonomer_52bp
TCCTCGGTCTTGGTATAAGTCGTCTATCGCATCCTATGAGTTACCGATCAAA

>HydrangeaTR13-consensusmonomer_89bp

### Supplementary Data

CTTGGAACATAAGGCGCCAAGGGGTGCAGTTCATTTCTGGTTTAGCATTGGAGCCTAAGGCATT
AAGACGCAGATCATTGCTATTGTTG

>HydrangeaTR14-consensusmonomer_185bp
GGAGGACGGTACAAATGCTCGTGTTCGGATTACGGTAAGACTAAAATAAACATTTGAAATTGGA
CTGTAAAACAAATTGAGATATCGAAAGAACTTTCCTTGTAAAGACGGAAATTATGAGTACCACA
TATGGAAATAAATTCCCACCAGTGGGATTTTTTTGAGTCTAAAATTGCAATTTTGTT

>HydrangeaTR15-consensusmonomer_846bp
CTAAGCTAGGTTCTGGCGCCGAGAACCTTCATGCACAGACTCACGTAGGCTCTGCAGGCAGTCA
CATGGTTTGTGATGCTTTTACCCGAGAATCCGGAAACTTGTCTCGGCATGACCCTAAGAGCATG
ATGCAAGTTACAAATATCCAAGGTAGTGCTTGCCACGTGGCAACAACCTCACCAAATGTTCAAA
TGCCAACCACTTTGTCATTACATGACCCTGCTGGCCATCTATTTACGCATACCTGCTTGAGTGG
TAATGAGGATGTTTGTACCACTAATCAAGAGAATGCTACCACCCCATCACAGCACATAACCACG
GCTCCTATTCCTAAGGCTGGAGTTGTTACAAAAGGTTCTTGCACAGCCCCACACCAGTCTGATA
CCCAAGTGAACCCTACCTACTGTGAGGCACTGAAGGCTAAATTACATAGTGATAACTCTGCTAC
GGCTAACTCCTCTGCAAAGCACATCCAAGAGACAAGCTGGGTTTCCCTATTGTAACCTAAAATA
GCTACAGAGTTTAAAGTCCATTATATCAAGCACCAGAGAGATATGGCCAGAATTCAAATTAGGG
TGCCTAGGATATTGACAGATCAAGGCAGCAAAGCTTGGGAGTACACTCTTGTGGGGCACTTCAT
AGGTAAGAAGCTTCCCTACTCTTTGGTGAAATCTGCATCCTCTAGACTCTGGGGTAGATTGGGA
CTTGAAGATATGTTAGCTATGGAATCAGGCTACTTCTTCTTCAAATTCAATTCCCAAGATGAAA
GTGCTGCCTCATGTTTTCCAGCTGGGAAAAGATCTGCAGCTGATCAGAACTTAGTTCTTGGCTT
GACGGCTTGCTCAG

#### Supplementary Data S2:

**Data S2: *Hydrangea*TR sequences used as FISH probes.** The alternating monomers are indicated by a change in color. Light markings indicate partial monomers.

>HydrangeaTR01-Hmac_207bp_OY742023.1
TCGTTATATTCTCAAAACGAGGGTTAACGCAAATCTGTTGCGAGAACCGAGAATATAAATCTGT
TATATTCCAAAACGGGGGTTAACGCAAATATGTTGCGAGAACCGAGAATATAAATCTGTTATAT
TCCAAAACGGGGGTTAACGCAAATCTGTTGCGAGAACCGAGAACATAAATCTGTTATATTCCAA
AAAGGGGGGTTAAAT

>HydrangeaTR03-Hmac_580bp_OY742025.1
GTATGTTTGAGGAGCGTGTAAGGTTTCGTGGCGTTTGGATAAGCGGTTTACCGATTAACTCCTC
AAAACCTCAATACTTAAGCTTTTTCAAGCATAAGTGTCTAAAGAGGCACTTTTTCGCATTTTTT
CGCATTTTTTTTCAGCCGTGAATCCGTAAAGCATTCCGTTAGTTCCCACAAGTTTATTTAGGTA
TGTTTGAGGTGCGTGTAAGGTTTCGTGGCGTTTGGTTAAGCGGTTTACCGATTAACTGCTCAAA
ACCTCAATACTTAAGCTTTTTCAAGCATAAGTGTCTAAAGAGCCACTTTTTCGCATTTTTTTTG
AACCGTGAATCCGTAAAGGATTCCGTTAGTTTACGCAAGTTTATGTAGGTATGTTTGAGGTGCG
TGTAAGGTTTCGTGGCATTTGGTTAAGCGGTTTACCGATTCACTACTCAAAACCTCAATACTTA

#### Supplementary Data

AGCTTTTTCAAGCATAAGTGTCTAAAGAGGCACTTTTTCGCATTTTTTTTGAACCGTGAATCCG
TAAAGCATTCCGTTAGTTCCCGCAAGTTTATTTAGGTATGTTAGAGGTGCGTGTAAGGTTTCGT
GGCG

>HydrangeaTR05a-Hmac_448bp_OY742027.1
CAAACCGTCGATCCGGTAACCACGAAACTTAATTCGCACCTCCCACACGTCCATAGAAAGTAAT
AAGAAGAAACAGAACAGGAATTGGACTCCCGGTTACCAAGATACAATATAAATCAACTTTTTTA
CCTCATAAACTCTTAAAAACTAAGTTTTTGGGGTTTGGGGGGTCAAAATTCAAACCGTCGATTC
GGTAACCACGAAACATAACTCGTGCTTCCCACACGTCCATAGAAAGTTGTAGGAAGAAACGGAA
CAGGAATTGGACTCCCGGTAATCAAGATACAATACAAAACAACTTTTTCACCTTATAGACTATG
AAAAATTAAGTTTTTGGGGTTTCGGAGATCAAAATGCAAACCGTCGATCTGGTAACCACGAAAC
TTAAGTCTCACCTCCCACAAATATAAAGAAAGTTGTAGGAAGAAACGGAACGGGAATCGGACTC
